# CcpNmr AnalysisDynamics: a unified framework for NMR dynamics data analysis

**DOI:** 10.64898/2026.06.19.733360

**Authors:** Luca G. Mureddu, Edward J. Brooksbank, Geerten W. Vuister, Frederick W. Muskett

**Author notes:** To whom correspondence should be addressed at or.

## Abstract

Nuclear Magnetic Resonance (NMR) relaxation experiments provide a powerful residue-resolved access to biomolecular dynamics across a wide range of timescales. Unfortunately, the quantitative analysis of the relaxation data remains distributed across specialised and often disconnected tools. Here, we present CcpNmr AnalysisDynamics, the latest addition to the CcpNmr Analysis program suite, providing an integrated framework for relaxation analysis, exchange-aware interpretation and dynamical modelling. The platform unifies relaxation-rate extraction, diagnostic validation, model-based analysis and structural visualisation within reproducible workflows, while supporting future extension through a robust application programming interface and plugin architecture. We introduce ModelAnalysis (ModA), a new analysis engine based on the Lipari-Szabo formalism that incorporates robust optimisation, uncertainty estimation and model-selection strategies designed for heterogeneous relaxation datasets. The framework also supports exchange-focused analysis and integration with specialised external modelling tools, allowing relaxation anomalies to be followed from initial detection to more detailed interpretation. The applicability and reliability of AnalysisDynamics are demonstrated through systematic re-analysis and validation of curated relaxation datasets from the Biological Magnetic Resonance Data Bank. These analyses enable assessment of data consistency, dynamic parameters and model reliability across magnetic fields, providing a reproducible route from NMR relaxation measurements to structure-linked interpretation of biomolecular dynamics.

## Introduction

Biomolecules and biomolecular complexes are intrinsically dynamic systems^1^. Their biological functions often crucially depend on motions spanning many orders of magnitude in time, from fast local fluctuations of atoms and side-chain rotations to slower conformational transitions, molecular recognition, assembly, folding and allosteric regulation^2^. Characterising these motions is therefore essential for understanding how molecular structure gives rise to biological function^1^.

Nuclear Magnetic Resonance (NMR) spectroscopy provides a uniquely powerful approach because it can report on dynamics with atomic or residue-level resolution across a broad temporal range. In biomolecular NMR, different experiments probe different regions of the dynamic timescale landscape (Fig. 1a). Fast picosecond–nanosecond motions, which report on local flexibility and internal mobility, are commonly characterised using longitudinal and transverse relaxation rates together with ^15^N heteronuclear NOE (hetNOE) measurements^3^. Motions on the microsecond–millisecond timescale, often associated with conformational exchange, enzyme motions and allosteric transitions, are probed using relaxation-dispersion approaches such as CPMG and R₁ρ experiments^4,5^. Slower millisecond–second processes, including folding, ligand binding and exchange between long-lived conformational states, can be accessed using methods such as CEST^6^, DEST^7^, ZZ-exchange^8^ and EXSY^9^. These experimental regimes are connected conceptually, but they require distinct forms of experimentation, data processing, validation and modelling^5^ (Supplementary Table 5).

**Figure 1.**
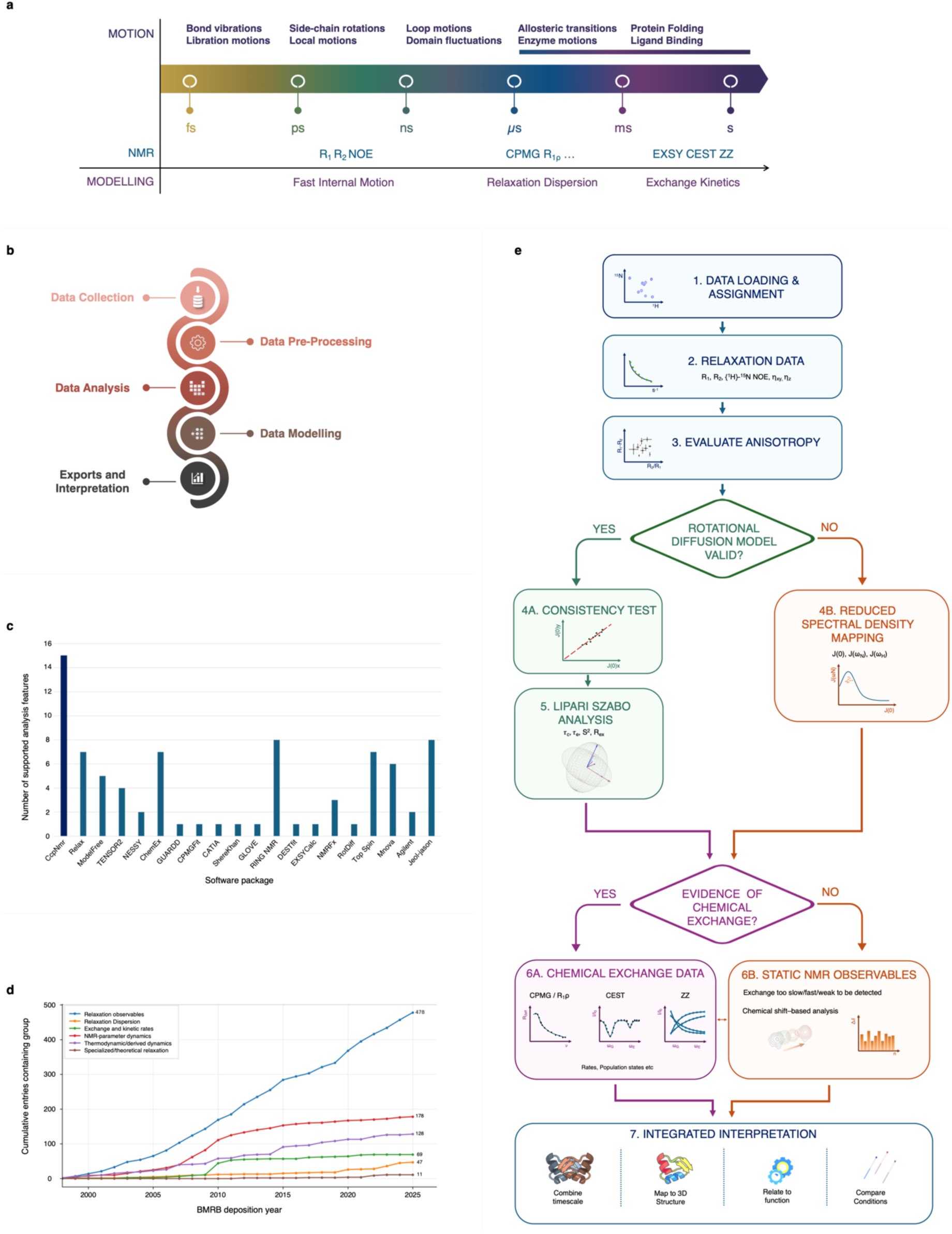
Conceptual landscape and workflow for biomolecular NMR dynamics analysis. **a,** Biomolecular motions occur over timescales ranging from femtoseconds to seconds. Different NMR experiments access different regions of this landscape, from R1, R2 and heteronuclear NOE measurements for fast picosecond–nanosecond dynamics to CPMG, R1ρ, CEST, DEST, ZZ-exchange and EXSY experiments for conformational exchange and kinetic processes. **b,** Schematic representation of the major stages of an NMR dynamics study, including data collection, preprocessing, analysis, modelling, export and interpretation. **c,** Feature coverage across representative NMR dynamics software packages, highlighting the fragmented support for different analysis and modelling tasks. **d,** Cumulative number of BMRB entries by class of NMR dynamics data, illustrating the growing availability of relaxation and exchange datasets for systematic reuse. **e,** Schematic AnalysisDynamics workflow for moving from assigned relaxation and exchange measurements through diagnostic validation, model-based analysis, exchange-focused analysis and structural interpretation.

A complete NMR dynamics analysis therefore extends well beyond the acquisition of relaxation or exchange data. It typically involves data collection, pre-processing, quantitative analysis, model selection, parameter estimation and structural or functional interpretation (Fig. 1b)^10,11^. In practice, however, analyses at these different stages are often handled by separate programs or involve user-written scripts or code. Existing tools tend to focus on specific experiment classes, particular theoretical models, or individual stages of the workflow (Supplementary Tables 6-7). This practice has produced a fragmented software landscape in which comprehensive dynamics studies frequently require semi-manual or ad hoc transfer of data between independent packages, each with its own file formats, assumptions and fitting conventions^12–15^.

This fragmentation is evident when the current NMR dynamics software ecosystem is compared across supported analysis features (Fig. 1c). While several mature programs^12–16^ provide robust implementations for specific tasks, few cover the full breadth of relaxation analysis, such as reduced spectral-density mapping, model-free analysis, relaxation dispersion, chemical-exchange kinetics and integrated visualisation^12,15^. As a result, combining complementary experiments that probe molecules across timescales remains technically demanding. The problem is particularly limiting for less routinely used approaches, including CEST, DEST and ZZ-exchange measurements, where analysis workflows are often less standardised and less accessible to non-specialist users^17^.

Beyond feature coverage, many existing workflows remain poorly connected to modern data management, graphical inspection of experimental data and accessible cross-platform execution^2,10,18^. As a result, it remains difficult to combine multiple experiment types within a single analysis environment and trace fitted parameters back to the underlying spectral and peak-level data. This need is reinforced by the growing number of publicly deposited NMR dynamics datasets, with the BMRB now containing multiple classes of relaxation and exchange experiments available for reuse, comparison and method development (Fig. 1d)^19^.

To address these limitations, we introduce CcpNmr AnalysisDynamics, a unified and extensible software resource for biomolecular NMR dynamics analysis. AnalysisDynamics integrates data analysis with dedicated higher-level modelling workflows for fast internal motion, relaxation dispersion and chemical exchange kinetics within a single coherent CcpNmr framework^20^. The software is designed to operate downstream of data collection and pre-processing, focusing on standardised analysis, modelling and interpretation steps, while maintaining direct links between quantitative outputs and the underlying spectral and peak-level data through the broader CcpNmr Analysis environment. This integration supports practical validation of results, error estimates and outlier assessment of fitted parameters without requiring users to move manually between disconnected tools.

AnalysisDynamics supports a broad range of established NMR dynamics experiments and modelling strategies and enables consistent parameter extraction across timescales. By combining these capabilities with cross-platform installation, an accessible graphical interface, documentation and tutorial-based workflows, the platform is intended to lower technical barriers to rigorous dynamics analysis within a best-practice framework. Here, we describe the design and implementation of AnalysisDynamics and demonstrate its application through representative use cases spanning relaxation analysis, diagnostic validation, model-based interpretation, exchange-focused workflows and systematic reanalysis of remediated public datasets.

## Results

We developed AnalysisDynamics to provide a flexible, high-level workflow for analysing biomolecular NMR dynamics from relaxation measurements to structural interpretation (Fig. 1e). The workflow is intended as a general framework that can be adapted to different molecular systems, experimental designs and biological questions, rather than as a single prescriptive analysis route.

The workflow typically assumes that the assignments of individual resonances of the molecule are known, e.g., ^1^H^N^ and ^15^N^H^, as it then allows for atom or residue-specific mappings. However, the program is not dependent on such information and can operate using any kind of labels, including enumerated numeric ones. The analysis can start from ^15^N-based relaxation datasets, including R_1_, R_2_ and hetNOE measurements and can incorporate additional diagnostic observables such as η_xy_ and η_z_ cross-correlated relaxation measurements^17,21^ for first-pass assessment of excess transverse relaxation. The workflow supports rate extraction, quality control, reduced spectral-density mapping, model-based analysis, exchange screening and visualisation of the resulting dynamic parameters.

A key feature of the workflow is the placement of diagnostic steps before model-dependent interpretation. Reduced spectral-density mapping is used to estimate J(0), J(ω_N_) and J(ω_H_), enabling residue-level inspection of relaxation behaviour and assessment of multi-field consistency before global fitting^22^. In parallel, η_xy_ and η_z_ analyses provide a practical screening layer by estimating the exchange-free transverse relaxation baseline, R_2,0_, from which apparent R_ex_ contributions can be inferred indirectly^21,23^. These measurements do not replace dedicated exchange experiments, but they provide residue-specific indicators that can be used to prioritise more detailed analyses.

Following these checks, the workflow directs the analysis either towards Lipari-Szabo model-free analysis^24^, when the relaxation data are compatible with the selected rotational diffusion model and no significant exchange contribution is detected, or towards exchange-focused analyses when additional chemical-exchange contributions are indicated. Candidate exchange sites identified from η_xy_- η_z_ -derived indicators, reduced spectral-density mapping, linewidth analysis or relaxation inconsistencies can then be followed up using exchange-sensitive experiments, including CPMG^25^, R_1ρ_^26^, CEST^16^ and ZZ-exchange^8^, within the same data analysis framework implemented in CcpNmr AnalysisDynamics. The resulting dynamic parameters can be mapped onto three-dimensional structure and compared across states or experimental conditions.

### Implementation of CcpNmr AnalysisDynamics

We implemented the AnalysisDynamics workflow as a modular backend framework within CcpNmr Analysis, exposed to users through an integrated graphical interface (Fig. 2). The tools operate directly on spectra, assignments and molecular data already present in the project, preserving the link between experimental measurements and derived dynamical parameters (Fig. 2a). This design minimises manual file conversion and enables relaxation, spectral-density and exchange analyses to be performed within a single reproducible framework.

**Figure 2.**
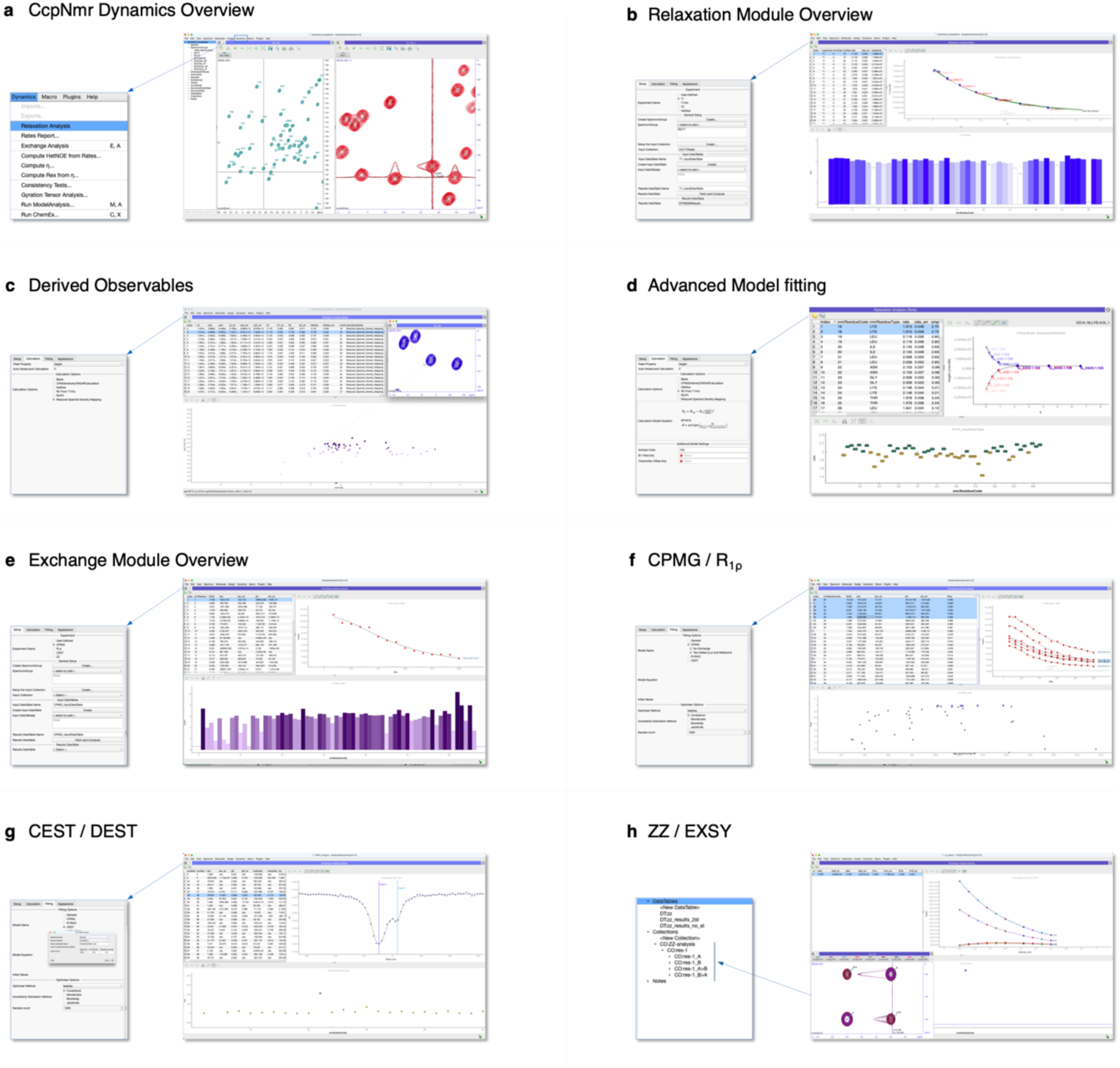
CcpNmr AnalysisDynamics overview and feature showcase. **a,** AnalysisDynamics extends the CcpNmr Analysis environment through dedicated dynamics menus and graphical interfaces, providing access to the main analysis, exchange and modelling modules. **b,** Relaxation module for extracting R1, R2 and heteronuclear NOE values from assigned spectral series and inspecting fitted curves, uncertainties and diagnostic plots. **c,** Derived-observables tools for reduced spectral-density mapping and other relaxation-based calculations that can be computed residue-wise and linked to the underlying data. **d,** Advanced model-fitting interface for shared or multi-dataset fitting and comparison of fitted parameters and diagnostics. **e,** Exchange module for computing exchange-derived parameters and organising exchange-sensitive datasets within the same project structure. **f,** CPMG and R1ρ interface for relaxation-dispersion analysis. **g,** CEST and DEST interface for saturation-transfer analysis. **h,** ZZ-exchange and EXSY interface for slower exchange and kinetic analysis.

The relaxation module provides dedicated routines for extracting R_1_, R_2_ and hetNOE values from series of assigned spectra (Fig. 2b). Fitted rates, uncertainties and diagnostic plots are generated alongside the original spectral data, allowing users to inspect both the primary measurements and the quality of the fit. Derived quantities are then passed directly to reduced spectral-density mapping and additional relaxation-based calculations, including residue-wise visualisation and comparison across datasets (Figs 2c, d). These tools make intermediate validation steps part of the standard analysis workflow, rather than separate post-processing procedures.

AnalysisDynamics also provides a unified interface for exchange-sensitive experiments. CPMG, R_1ρ_, CEST and ZZ-exchange datasets can be organised, fitted and inspected using the same project structure used for fast-timescale relaxation analysis (Figs 2e-h). This allows exchange data to be interpreted together with R_1_, R_2_, hetNOE and spectral-density outputs, supporting analysis across complementary motional regimes. Because these analyses remain linked to the same spectra, assignments and molecular identifiers, users can trace exchange-derived parameters back to the experimental observations from the same system from which they were obtained.

### The ModA and ChemEx interfaces

For advanced modelling, AnalysisDynamics provides graphical wrappers for specialised analysis engines (Fig. 3). The ModA interface supports Lipari-Szabo model analysis of fast-timescale backbone dynamics, whereas the ChemEx^16,27^ interface enables preparation, execution and reintegration of chemical-exchange analyses. In both cases, input generation and result parsing are handled inside the CcpNmr environment, allowing specialised algorithms to be run in parallel processing without breaking the connection to the underlying project.

**Figure 3.**
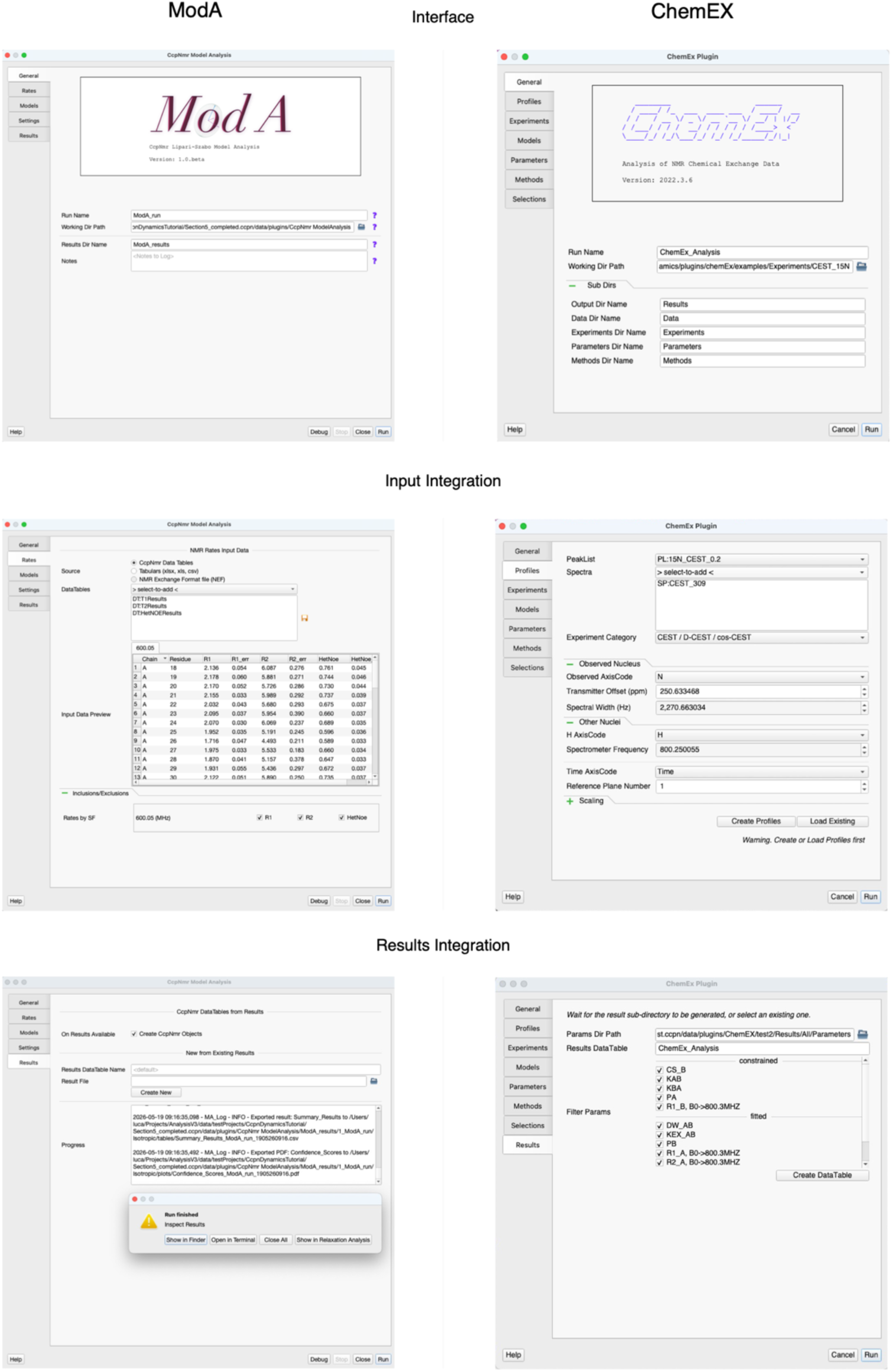
ModA and ChemEx interfaces for advanced dynamics modelling. AnalysisDynamics provides graphical plugin interfaces for advanced modelling workflows, illustrated here for ModA and ChemEx. ModA is integrated as a native AnalysisDynamics modelling workflow for Lipari-Szabo analysis from relaxation inputs, whereas the ChemEx plugin provides a graphical wrapper for preparing, running and reintegrating analyses performed with the external ChemEx package. Both interfaces support input preparation, calculation setup and structured result reintegration within the CcpNmr Analysis environment, preserving links between fitted parameters, diagnostics, spectra and assignments.

ModA fits residue-resolved relaxation observables and associated uncertainties to a hierarchy of spectral-density parameterisations and returns dynamic parameters, uncertainty estimates and model-selection metrics. A distinguishing feature of ModA is its adaptive per-residue bound definition for τ_e_, which uses local relaxation behaviour and the estimated global tumbling time to define residue-specific search ranges for internal correlation times. ModA also uses parallel residue-wise execution by default, allowing independent model fits and Monte Carlo refits to be distributed across available worker processes. The full implementation, including supported models, fitting options, parameter-initialisation procedures, algorithmic details and workflow features, is described in Supplementary Table 8 and Supplementary Note 3.

To evaluate numerical agreement with established implementations, we compared ModA-derived order parameters with those obtained using RotDif version 3.1^13^, ModelFree version 4^14^ and Relax version 5^12^ for the same ubiquitin benchmark dataset under an isotropic rotational-diffusion model (Fig. 4a). Pairwise S² similarity scores between packages were high overall (Fig. 4b-c), ranging from 0.95 to 0.98. ModA showed strong agreement with ModelFree and Relax, with similarity scores of 0.96 and 0.95, respectively. Residue-wise deviation analysis showed that ModA reproduced the established order-parameter landscape, while also revealing modest implementation-dependent differences between packages. Because all calculations used an isotropic diffusion description, these differences are unlikely to reflect gross differences in the tumbling model. Instead, they are more likely to arise from package-specific settings and conventions, such as the ^15^N chemical-shift anisotropy, N-H bond length, optimisation tolerances, parameter bounds and model-selection procedures.

**Figure 4.**
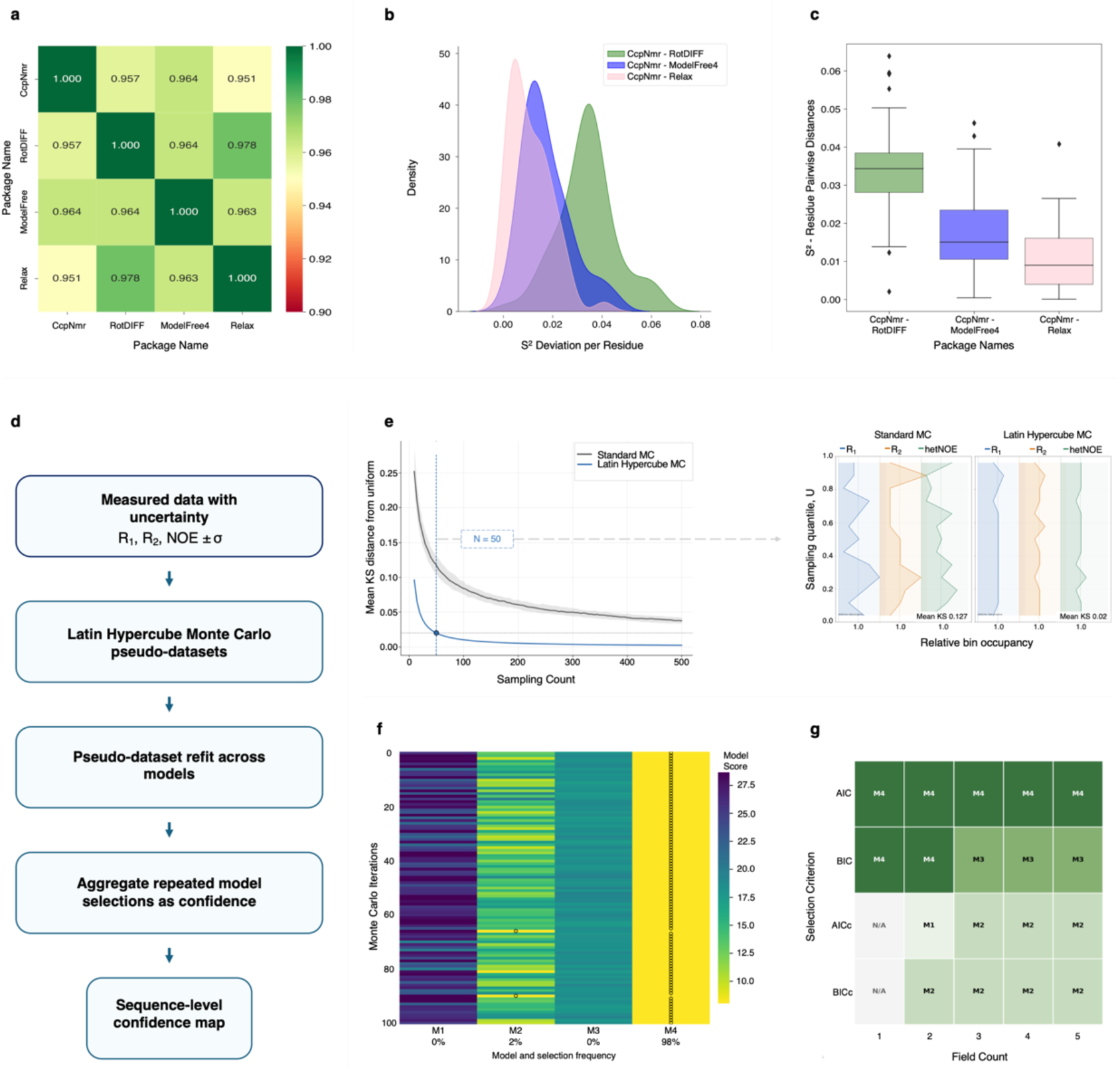
Benchmarking and confidence scoring in ModA. **a,** Pairwise similarity matrix of residue-level Lipari-Szabo order parameters (S²) calculated with ModA/CcpNmr, RotDif, ModelFree and Relax using the same ubiquitin benchmark dataset. **b,** Density distributions of residue-wise S² deviations between ModA/CcpNmr and each comparator package. **c,** Box-plot analysis of the same residue-wise deviations, summarising the median, dispersion and outliers for each comparison. **d,** Model-selection confidence scoring by uncertainty propagation. The schematic summarises the analysis procedure: measured R1, R2 and hetNOE uncertainties are used to generate Latin Hypercube Monte Carlo pseudo-datasets, each pseudo-dataset is refitted across candidate models and repeated model selections are aggregated into residue-level and sequence-level confidence scores. **e,** Sampling-convergence and bin-occupancy analyses comparing coverage efficiency between standard Monte Carlo and Latin Hypercube sampling. Latin Hypercube sampling provides more uniform sampling of the uncertainty space than standard Monte Carlo sampling. **f,** Residue-level heat map showing iteration-wise model scores and model-selection frequency for an example residue. **g,** Field-count matrix showing how the selected model changes with the number of fitted magnetic-field datasets and with the information criterion used for model selection.

We next assessed how uncertainty calibration affects χ²-based fit diagnostics before model selection. Synthetic replicas of the ubiquitin relaxation dataset were generated by scaling the relative experimental uncertainties of R_1_, R_2_ and hetNOE observables over several orders of magnitude while preserving their observable-specific error structure (Supplementary Fig. 1a). This distinction is important because relaxation rates and hetNOE measurements can exhibit different noise behaviour, with R_1_ and R_2_ errors often approximated as additive and hetNOE-derived quantities being more sensitive to relative or multiplicative error. Full details of the uncertainty-scaling procedure, replica generation and balanced χ² algorithm are provided in Supplementary Note 3.

For each uncertainty-scaled dataset, ModA fitted the relaxation data using a fixed model configuration. This configuration was used only as a controlled baseline for testing χ² sensitivity to input uncertainty, not for model selection or physical interpretation. The final χ² value was calculated for each residue, and the median residue-level χ² was evaluated as a function of the imposed fractional relative error (Supplementary Fig. 1b). Standard χ² depended strongly on the assigned uncertainty scale, whereas the balanced χ² remained close to numerical precision across the scaling series. These results show that uncertainty calibration directly affects χ²-based fit assessment and should be considered before downstream model selection.

To propagate experimental uncertainty into model selection, ModA generates Monte Carlo pseudo-datasets from the measured R_1_, R_2_ and hetNOE uncertainties, refits each pseudo-dataset across the candidate Lipari-Szabo model hierarchy and aggregates repeated model selections into confidence scores (Fig. 4d). Because repeated fitting is computationally expensive, we evaluated standard Monte Carlo and Latin Hypercube Monte Carlo sampling in a three-observable uncertainty space. Latin Hypercube sampling converged more rapidly towards uniform quantile coverage, giving a mean Kolmogorov-Smirnov distance of 0.020 at N = 50 compared with 0.127 for standard Monte Carlo. Bin-occupancy profiles confirmed more even marginal coverage across R_1_, R_2_ and hetNOE uncertainty dimensions (Fig. 4e).

The resulting model-selection frequencies provide a direct residue-level confidence measure. In the example shown, M4 was selected in 98% of uncertainty replicas, whereas M2 was selected in 2% and M1 and M3 were not selected, indicating a robust model assignment for that residue (Fig. 4f). The field-count analysis further shows that model selection depends on both the amount of experimental information and the information criterion used. In the simulated example, AIC favoured the most complex model across all field counts, whereas BIC and corrected criteria selected more parsimonious models as the number of fitted datasets changed (Fig. 4g). These diagnostics allow ModA to report not only a selected model, but also the robustness and criterion-dependence of that assignment.

### ModA validation of BMRB datasets

Relaxation measurements collected at multiple magnetic fields must be internally consistent before they are combined in quantitative dynamics analyses^28^. AnalysisDynamics therefore estimates J(0) independently at each field and compares field-specific distributions using standard graphical outputs, including J(0)-J(0) comparisons, quantile-quantile plots and PCA-derived diagnostics. Field-dependent deviations can indicate experimental bias, sample-condition differences or problematic measurements, but may also reflect genuine exchange contributions or other residue-specific dynamics; flagged datasets therefore require inspection rather than automatic exclusion.

We applied these diagnostics to 70 curated relaxation datasets from the BMRB. Of these, 37 contained relaxation measurements at multiple magnetic fields and were thus suitable for consistency testing; an internal GB1 dataset was included as a positive control. For this comparative survey, the individual diagnostics were normalised across datasets and combined into an aggregate inconsistency score to rank field-to-field agreement. This aggregate score was used only for the BMRB-level analysis, whereas the underlying per-metric plots and diagnostics are available as standard AnalysisDynamics outputs.

The radial plot (Fig. 5a) shows the contribution of individual metrics to the aggregate score, indicating that inconsistency can arise from different sources, including distributional shifts, residue-wise fractional changes or multivariate outliers. Across the 37 multi-field datasets, the total normalised inconsistency score ranged from 0.036 for GB1 to 0.709 for the most inconsistent dataset. Based on these scores, 34% of datasets were classified as good, 32% as average, 32% as poor and 3% as very poor.

**Figure 5.**
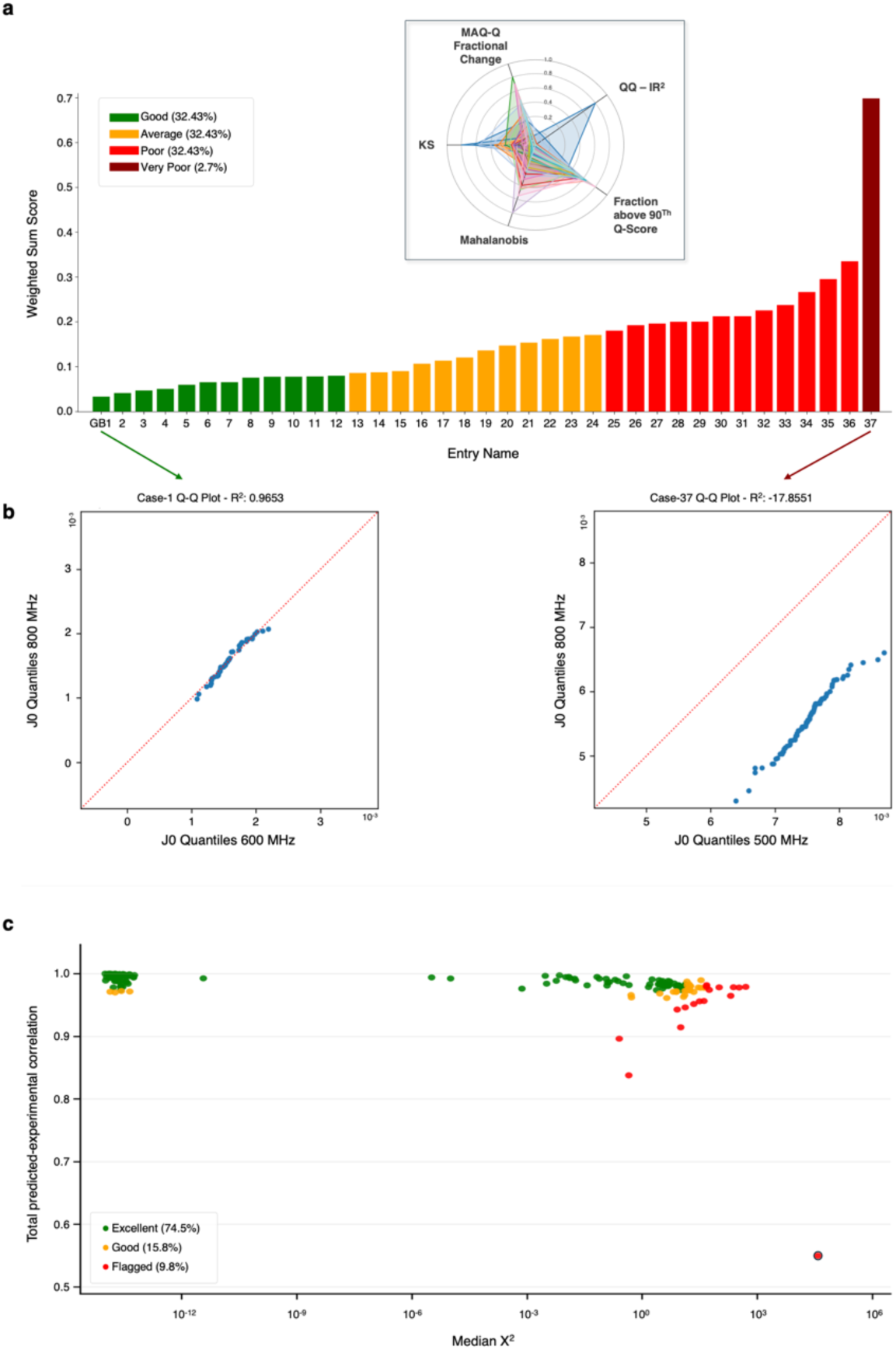
Consistency testing and empirical fit quality across curated relaxation datasets. **a,** Multi-field consistency analysis of relaxation datasets. The radial plot shows the contribution of individual diagnostics to the survey-level inconsistency score, including KS, Q-Q agreement, fractional-change, Mahalanobis and PCA-derived Q-score metrics. The bar plot ranks the 37 multi-field datasets by aggregate normalised inconsistency score, with lower values indicating stronger agreement across fields. **b,** Example Q-Q plots show a consistent GB1 control dataset with J(0) quantiles close to the diagonal (left) and the most inconsistent dataset, with strong deviation from the diagonal (right). **c,** Empirical ModA fit quality across all analysed field combinations. Each point represents one ModA run, assessed by total experimental-predicted Spearman correlation and median residue-level χ². Runs were classified as excellent, good or flagged, with most analyses showing high agreement between experimental and back-calculated relaxation observables.

The representative quantile-quantile plots (Fig. 5b) illustrate the extremes of this distribution. For GB1, J(0) quantiles remained close to the diagonal, yielding a Q-Q R² value of 0.96. In contrast, the most inconsistent dataset deviated strongly from the diagonal, with a Q-Q R² value of -17.85. These examples show that automated consistency testing provides a practical quality-control layer before further modelling, while retaining the graphical diagnostics needed to assess whether poor agreement reflects technical inconsistency, residue-specific outliers or behaviour requiring further inspection.

We next analysed the curated relaxation datasets with ModA across all available magnetic-field combinations. For datasets recorded at multiple fields, ModA was run on individual fields and on all pairwise or higher-order field combinations, generating 184 total analyses across the 70 public datasets and two internal controls. Fit quality was assessed by comparing experimental relaxation observables with rates back-calculated from the selected Lipari-Szabo models, using the total Spearman correlation between observed and simulated rates together with the median residue-level χ² (Fig. 5c).

The analysis showed strong agreement between experimental and back-calculated relaxation rates. Overall, 75% of runs were classified as excellent, 16% as good and 10% as flagged. The high proportion of excellent and good fits indicates that ModA captured the dominant relaxation behaviour across a heterogeneous collection of datasets. Flagged cases were characterised by lower rate correlations and/or elevated median χ² values, consistent with problematic input data, field-specific inconsistencies or localised residue-level outliers rather than a general failure of the fitting procedure.

Inspection of flagged runs indicated that several poor scores could be traced to data-quality or metadata issues, including cases where relaxation observables were misassigned or incompletely represented in the deposited dataset. Other flagged cases may reflect older datasets acquired or processed before current standards for relaxation-data reporting and uncertainty estimation were widely adopted. These results show that ModA can be applied systematically across diverse archived datasets, while also providing quantitative diagnostics that identify cases requiring manual inspection before biological interpretation.

## Discussion

NMR relaxation experiments provide a uniquely detailed view of biomolecular dynamics^10,18^, but their practical analysis remains difficult because experimental fitting, data validation, Lipari-Szabo model-free analysis^24^, exchange analysis and structural interpretation are often performed using separate tools^12,16,24^. CcpNmr AnalysisDynamics addresses this fragmentation by placing these steps within a single project-based environment. By preserving the connection between spectra, assignments, fitted rates, derived quantities and final dynamic parameters, the platform reduces manual data transfer and provides a more reproducible route from experimental measurements to molecular interpretation.

A central principle of the AnalysisDynamics workflow is that model-dependent analysis should be preceded by explicit diagnostic testing (Fig. 1e). Reduced spectral-density mapping and multi-field consistency analysis provide an intermediate layer between relaxation-rate fitting and dynamical modelling. This diagnostic layer is necessary because relaxation datasets can appear suitable for Lipari-Szabo analysis^24^ while still containing field-dependent inconsistencies, outlying residues or exchange contributions that violate the assumptions of joint fitting. By incorporating these checks into the standard workflow, AnalysisDynamics shifts quality control from an optional post-processing step to a prerequisite for quantitative interpretation.

The implementation of ModA further extends this framework by providing a statistically controlled Lipari-Szabo analysis engine within AnalysisDynamics (Fig. 3). The benchmark against established software showed that ModA reproduces comparable order parameters, while the additional χ^2^ diagnostics and Monte Carlo model-selection framework provide information not captured by a single best-fit model. In particular, model-selection confidence scores help distinguish residues with robustly supported motional descriptions from residues for which the available relaxation data are insufficient to discriminate between competing models. This distinction is important for avoiding overinterpretation of fitted parameters, especially in heterogeneous or noise-limited datasets (Fig. 4).

The public BMRB analysis highlights a broader issue for the field: convergence of a model-free fit is not, by itself, evidence that the input data support a reliable physical interpretation. The consistency test analysis^22^ revealed substantial variability among multi-field relaxation datasets, indicating that some public datasets are well suited for joint analysis whereas others require filtering, field-specific inspection or more cautious interpretation. This result supports the placement of spectral-density consistency testing before Lipari-Szabo fitting and suggests that systematic re-analysis of public relaxation data can provide useful benchmarks for both software development and experimental practice.

Several limitations remain. As with all relaxation-based approaches, the reliability of fitted dynamic parameters depends on the quality of the input relaxation rates, the accuracy of reported uncertainties, the appropriateness of the rotational diffusion model and the number and diversity of experimental observables. J(0)-based consistency testing can identify field-dependent discrepancies, but cannot by itself determine whether these arise from experimental artefacts, sample differences, incomplete uncertainty estimates or genuine conformational exchange. Similarly, ModA provides uncertainty-aware model selection and confidence estimates, but these remain conditional on the candidate model hierarchy and the assumptions of the Lipari-Szabo formalism. These limitations reinforce the need for workflows in which diagnostic testing, model fitting and biological interpretation are not treated as independent steps, but as linked components of a single analysis process.

A further strength of AnalysisDynamics is that it does not restrict dynamics analysis to fast-timescale relaxation or to a single model-free interpretation. The exchange-analysis layer provides native support for organising and analysing CPMG^29^, R_1ρ_^26^, CEST^16^ and ZZ-exchange experiments within the same project framework used for R_1_, R_2_, hetNOE and spectral-density analysis (Fig. 2). By linking exchange-sensitive datasets to the same spectra, assignments, residue identifiers and structural models, AnalysisDynamics allows complementary relaxation and exchange experiments to be interpreted within a common data environment.

The integration of ChemEx^27^ extends this capability further by connecting AnalysisDynamics to specialised exchange-modelling algorithms without requiring users to leave the CcpNmr project environment. Input files can be generated from structured AnalysisDynamics objects, external calculations can be executed through the plugin interface and fitted exchange parameters can be parsed back into the project for inspection, comparison and structural mapping. This design preserves access to specialist modelling approaches while avoiding the loss of provenance that can occur when data are repeatedly exported, reformatted and analysed in separate programs. In this respect, ChemEx integration is not simply an additional fitting option, but a demonstration of how AnalysisDynamics can act as a coordinating environment for advanced dynamics analysis.

The underlying plugin framework is therefore central to the long-term value of the platform. Native modules and external engines are represented through a common execution and data-handling model, in which input generation, process execution, result import and downstream visualisation are handled consistently. Registered namespaces, collection structures and observable-context trees allow simple relaxation series, multi-field datasets, state-specific observables and exchange pathways to coexist within the same project while remaining distinguishable and reproducible. This architecture also provides a route for future extensions, including additional relaxation observables, new exchange models, alternative optimisation engines, validation metrics and reporting tools.

## Conclusions

In summary, CcpNmr AnalysisDynamics was developed to make biomolecular NMR dynamics analysis more reliable, reproducible and directly linked to structural and functional interpretation. Dynamics measurements are central to understanding how biomolecules fluctuate, exchange between conformational states and distribute mobility across functional sites, but these interpretations depend on careful handling of relaxation data, uncertainties, model assumptions and structural context. AnalysisDynamics addresses this need by connecting the full analysis path within a single CcpNmr Analysis project, from spectra and assignments to fitted rates, diagnostic tests, model parameters, exchange analyses and structure-mapped outputs. This integrated framework also facilitates the routine use of experiments that provide valuable insight into biomolecular dynamics but have often been underused because of the practical complexity of their analysis.

The scientific value of AnalysisDynamics lies in its validation-first approach to dynamics interpretation. Reduced spectral-density mapping, multi-field consistency testing and exchange diagnostics help identify when relaxation data support Lipari-Szabo analysis, or when further inspection is needed. Within this framework, ModA provides a new implementation of Lipari-Szabo analysis that extends conventional model-free workflows with uncertainty-aware fitting, model-selection confidence scoring and integrated diagnostic assessment.

AnalysisDynamics also supports dynamics analysis beyond fast-timescale relaxation. Native tools for CPMG, R_1ρ_, CEST and ZZ-exchange, together with ChemEx integration, allow exchange-sensitive datasets and specialised modelling outputs to be handled within the same project structure. More broadly, a new plugin architecture provides a route for adding new observables, models, external engines and reporting tools, making AnalysisDynamics both accessible for routine use and adaptable for future methods development.

## Online Methods

### Software architecture, execution model and plugin framework

CcpNmr AnalysisDynamics is implemented as a new application within the CcpNmr Analysis suite, providing workflows for NMR relaxation analysis, exchange analysis and dynamical modelling. Modular backend APIs are exposed through graphical interfaces and operate directly on spectra, assignments, molecular information and derived data stored within a CcpNmr project. Intermediate and final outputs are written to structured project objects, preserving links between input data, fitted parameters, uncertainties, model metadata and downstream visualisation.

Native modules and external analysis engines are supported through a plugin framework. External engines are executed as independent operating-system processes, while execution state, logs, return codes and output files are monitored by the host application. Results are parsed, standardised and reintegrated into the AnalysisDynamics data model for validation, visualisation and reporting. Additional implementation details are provided in Supplementary Notes 1 and 2.

### Relaxation and exchange analysis workflows

Relaxation and exchange analyses are implemented as so-called CcpNmr series-based workflows in which spectral observables are measured across ordered experimental variables such as relaxation delays, saturation states, field strengths, mixing times or offsets. Input data are generated from CcpNmr spectrum groups and peak collections, then stored as structured data tables together with model metadata, fitted parameters, uncertainties, fitting statistics and exclusion flags. Calculation models perform deterministic transformations, such as hetNOE calculation or conversion of intensity series to exchange observables, whereas fitting models estimate parameters using analytical or numerical equations.

For standard relaxation analysis, the framework supports extraction, inspection and curation of R_1_, R_2_ and hetNOE values, including weighted fitting, residue-wise refitting, multi-residue global fitting and propagation to reduced spectral-density mapping and ModA analysis. Exchange experiments are handled by a dedicated analysis layer because CPMG, R_1ρ_, CEST and ZZ-exchange data require experiment-specific pre-processing, multiple candidate models, context-dependent parameters and in some cases, custom objective functions or staged fitting pipelines. CcpNmr Analysis DataTable objects are resolved using registered namespaces and observable-context collection trees, enabling simple relaxation series and state- or path-specific exchange observables to be represented within the same project. Details are provided in Supplementary Note 1 and Supplementary Fig. 2.

### ModA relaxation modelling framework

ModA interprets heteronuclear relaxation observables using the Lipari-Szabo spectral-density formalism^24^. Experimental R_1_, R_2_ and steady-state hetNOE values, together with their uncertainties, are fitted by forward calculation of relaxation rates from candidate spectral-density parameterisations. The implementation separates spectral-density models, rotational diffusion models, relaxation-rate equations and numerical optimisation, allowing the same model hierarchy to be evaluated consistently across observables, magnetic fields and rotational diffusion descriptions.

For each residue, ModA evaluates a hierarchy of Lipari-Szabo spectral-density parameterisations with different combinations of order parameters, effective internal correlation times and optional exchange contributions to R_2_. Overall rotational diffusion is represented using isotropic, axially symmetric or fully anisotropic models. Structural information is used only when anisotropic diffusion models require N-H bond-vector orientations relative to the diffusion frame. Full spectral-density definitions, relaxation-rate calculations and diffusion-model equations are provided in Supplementary Note 3.

### Optimisation, uncertainty estimation and model selection

Dynamic parameters are estimated by minimising the difference between experimental- and back-calculated relaxation observables for each residue and candidate spectral-density model. Optimisation is performed under physically motivated parameter bounds using gradient-free global optimisation strategies suited to the nonlinear parameter landscapes encountered in Lipari-Szabo analysis. Initial estimates of global tumbling and local internal motion are used only to initialise the optimisation and do not constrain the final fitted parameters.

To reduce sensitivity to unreliable experimental uncertainties, ModA uses a noise-aware objective function that stabilises the weighting of different relaxation observables during fitting. Parameter uncertainties are estimated by Monte Carlo resampling of the experimental relaxation data. Pseudo-datasets are generated in uncertainty space and refitted using the same optimisation workflow, yielding parameter distributions from which uncertainty estimates are derived.

Candidate models are compared using likelihood-based information criteria, including small-sample-corrected forms where appropriate. Model-selection robustness is assessed by repeating model comparison across Monte Carlo pseudo-datasets. The model-selection confidence score is defined as the fraction of resampled datasets for which a given model is preferred, allowing residues with robust and ambiguous model assignments to be distinguished. Details of the optimisation strategy, objective-function weighting, Monte Carlo sampling and model-selection scoring are provided in Supplementary Note 3.

### Global tumbling refinement, structural integration and consistency testing

Before residue-level optimisation, ModA estimates the overall rotational correlation time using a modular procedure based on grouped R_1_ and R_2_ measurements at each spectrometer field. The default implementation follows the relaxation-rate ratio approach of Fushman and co-workers^30^, in which τ_c_ is estimated from longitudinal and transverse relaxation times under the assumption that overall tumbling dominates the relaxation behaviour. Residues with anomalous relaxation properties are filtered before this estimate is computed, reducing bias from flexible sites, exchange-affected residues or poorly determined rates. This initial τ_c_ is used only to initialise the diffusion parameters and is not interpreted as the final global tumbling estimate. After residue-level fitting, ModA optionally refines the global tumbling description by optimising a shared diffusion-scaling factor while keeping residue-specific internal parameters fixed. This final refinement enforces a consistent global description of overall tumbling across the fitted dataset.

For anisotropic diffusion models, atomic coordinates are obtained from standard structure files and used to define molecular-shape descriptors and backbone N-H bond-vector orientations relative to the diffusion frame. Structural information is used only to define rotational geometry and does not impose additional assumptions on internal motion.

Before joint fitting of relaxation data acquired at multiple magnetic fields, AnalysisDynamics assesses internal consistency using reduced spectral-density quantities calculated independently at each field. Residue-wise, distribution-level and multivariate comparisons are used to identify field-dependent discrepancies that could compromise joint fitting. These tests provide a quality-control step before ModA analysis. Details of global correlation-time estimation, final tumbling refinement, structural integration and consistency testing are provided in Supplementary Notes 3 and 4.

### Public datasets and remediation criteria

A set of 70 publicly deposited biomolecular NMR relaxation datasets was downloaded from the BMRB, curated and analysed with AnalysisDynamics using multiple analysis protocols. Datasets were retained when residue-resolved relaxation observables, magnetic-field information and sufficient metadata were available to reconstruct the analysis inputs. During remediation, entries were standardised for residue identifiers, observable type, relaxation rates, spectrometer frequencies and uncertainty values, with numerical quantities converted to SI units using automated remediation scripts. Multi-field datasets were further evaluated using the consistency-testing workflow described above. Remediated BMRB datasets are available at https://github.com/ccpnmr/curated-bmrb-relaxation-dynamics.

## Code availability

CcpNmr AnalysisDynamics is available to download from ccpn.ac.uk/downloads as part of the CcpNmr suite. Source code is available at https://github.com/ccpnmr/analysisdynamics.

## Acknowledgements

We thank Dr Victoria Higman for proofreading the manuscript and for valuable feedback on AnalysisDynamics workflows, software testing and usability of the graphical interface. We are grateful to the whole CcpNmr team for their support, technical discussions and feedback throughout the development, testing and refinement of AnalysisDynamics. We acknowledge the Biological Magnetic Resonance Data Bank for access to publicly deposited NMR relaxation datasets.

## Author contributions

**LGM:** Conceptualisation, Data Curation, Methodology, Software, Formal analysis, Investigation, Data curation, Visualisation, Writing - original draft, review and editing.

**EJB:** Software.

**GWV**: Conceptualisation, Supervision, Software, Funding acquisition, Writing - review and editing.

**FWM:** Data acquisition, Investigation, Methodology, Resources, Validation, Writing - original draft, review and editing.

## Abbreviations

AIC: Akaike information criterion
AICc: corrected Akaike information criterion
BIC: Bayesian information criterion
BICc: corrected Bayesian information criterion
BMRB: Biological Magnetic Resonance Data Bank
CEST: chemical exchange saturation transfer
CPMG: Carr-Purcell-Meiboom-Gill
DEST: dark-state exchange saturation transfer
EXSY: exchange spectroscopy
GB1: immunoglobulin-binding domain B1 of protein G
GUI: graphical user interface
hetNOE: ^15^N heteronuclear NOE
KS: Kolmogorov-Smirnov
LHS: Latin Hypercube sampling
MC: Monte Carlo
ModA: ModelAnalysis
NMR: nuclear magnetic resonance
PCA: principal component analysis
Q-Q: quantile-quantile
R_1_: longitudinal relaxation rate
R_1ρ_: rotating-frame longitudinal relaxation rate
R_2_: transverse relaxation rate
R_ex_: exchange contribution to transverse relaxation
RSDM: reduced spectral-density mapping
CCPN: Collaborative Computational Project for NMR
_S_2: Order Parameter
_χ_2: Chi-squared
ZZ: longitudinal two-spin order exchange experiment

