## Supplementary information for "CcpNmr AnalysisDynamics: a unified framework for NMR dynamics data analysis"

### Supplementary Note 1. Software architecture, execution model and plugins framework

CcpNmr AnalysisDynamics is implemented as a modular extension within the CcpNmr Analysis environment and builds upon its established data model, execution infrastructure and graphical framework[1,2]. The software is written in Python and uses the scientific computing libraries provided as part of the CcpNmr Analysis distribution. Core numerical operations are performed using NumPy[3,4], while optimisation routines are drawn from SciPy and related libraries[3,4]. Parameter handling, bounds management and refinement are implemented using LMFIT[5], which enables consistent definition and manipulation of parameter sets across different analysis modules. Experimental data, intermediate results and summary tables are organised using tabular data structures to support efficient manipulation and integration with downstream analysis and reporting.

The internal architecture of AnalysisDynamics is organised around a layered design that separates data representation, theoretical models, optimisation logic and result aggregation. Higher-level analysis engines, including the Lipari–Szabo–based ModelAnalysis (ModA) module, are implemented within this architecture and interact with the broader framework through well-defined interfaces, as described in more detail in Supplementary Note 3.

#### Relaxation Analysis Module

The Relaxation module is implemented within the general Series Analysis Framework of CcpNmr Analysis. This framework provides a common API for experiments in which a spectral observable is measured across an ordered experimental series, such as relaxation delays, saturation states or field-dependent conditions. Input data are generated directly from CcpNmr SpectrumGroup objects and associated peak Collection objects, preserving the connection between spectra, peak intensities, assignments and derived relaxation parameters (Figs 2a-c). Results are stored in CcpNmr DataTable objects together with model metadata, fitted parameters, uncertainties, fitting statistics and exclusion flags, allowing analyses to be saved, restored and propagated to downstream workflows.

The implementation separates deterministic calculation models from fitted models. Calculation models perform direct transformations of the input data, such as heteronuclear NOE (hetNOE) calculation from saturated and unsaturated spectra, with uncertainty propagation from peak signal-to-noise estimates. Fitting models estimate parameters from experimental series data using defined analytical equations, including exponential decay models for relaxation-rate extraction. Each model provides its equation, parameters, documentation and default settings and models are registered dynamically with the CcpNmr software so that new calculation or fitting routines can be added without modifying the core framework.

Relaxation-rate fitting is performed residue-wise by grouping series data according to peak collections or assignments. The Relaxation module supports weighted fitting using peak figure-of-merit values, configurable minimisation and uncertainty-estimation settings and refitting of individual residues with alternative fitting models when required. It also supports global fitting across multiple residues, in which selected parameters can be shared, local or fixed, enabling multi-residue fits with common physical parameters while preserving residue-specific terms.

The graphical interface exposes these functionalities through predefined experiment setups, model selection, result tables and customisable plots (Figs 2b-d). Fitted curves are regenerated from the selected model and fitted parameters, allowing direct inspection of the fits for individual residues or atoms. Exclusions can be applied to residues, peaks, atoms, collections or spectra and these are stored in the output table with the ability to reverse the decision. Excluded values are masked from downstream analyses while remaining traceable in the project. This design allows relaxation observables such as $R_1$, $R_2$ and hetNOE values to be extracted, inspected, curated and passed reproducibly to spectral-density mapping and ModA analysis.

**Other pipelines**

The same API also enables specialised relaxation workflows to be implemented as lightweight extensions rather than separate analysis programs. For example, dedicated graphical popups can orchestrate multi-step analyses such as $\eta_{xy}/\eta_z$ extraction from HSQC- or TROSY-based experiments[6,7], or Rex calculation from previously fitted $R_1$, $R_2$ and η-derived data tables. These workflows reuse the same backend objects, input table construction, calculation-model registry, fitting-model registry and output DataTable infrastructure as the standard relaxation module. As a result, new experiment-specific protocols can combine multiple SpectrumGroup inputs, generate peak collections, create intermediate data tables, apply the appropriate calculation and fitting models and write results back into the CcpNmr project without duplicating the core analysis logic. This design allows AnalysisDynamics to support both predefined user-friendly workflows and advanced custom protocols while maintaining consistent data provenance, model metadata and downstream compatibility.

**Exchange Analysis Module**

The Exchange Analysis module is implemented as a separate analysis layer because NMR exchange experiments require more heterogeneous data handling and model fitting compared to standard relaxation-rate analysis (Fig. 2d). Whereas $R_1$, $R_2$ and hetNOE analyses usually involve one experimental series and a small number of fitting models, exchange experiments often require experiment-specific preprocessing, multiple independent variables, multiple input tables, context-dependent parameters and several competing physical models[8]. For example, CPMG relaxation-dispersion data may be analysed using no-exchange models, analytical two-state approximations such as Luz-Meiboom, Carver-Richards or Ishima-Torchia models, or numerical Bloch-McConnell models (Fig. 2f)[9,10]. Related requirements arise for $R_{1\rho}$, $R_{2\rho}$, CEST and ZZ-exchange experiments, each of which has distinct observables, acquisition variables and model assumptions (Fig. 2g-h)[11–13].

The Exchange Analysis module therefore uses a dedicated exchange-domain fitting API. Input data from one or more DataTable objects are normalised, merged and grouped into fit units, typically by residue or assignment. Each fitting model then constructs a standard fit payload containing the observed values, optional weights, independent variables, experimental contexts and model-specific metadata. This abstraction allows the same fitting framework to support simple one-dimensional dispersion curves, multi-field datasets with context-local baselines, array-valued CEST profiles and other future exchange experiments with any number of control variables.

Exchange fitting models are separated into model families, including CPMG, $R_{1\rho}$, CEST and ZZ exchange. Models may be analytical, using closed-form approximations to the Bloch-McConnell equations, or numerical, in which the magnetisation evolution is simulated directly. Analytical models provide rapid screening and comparison when their assumptions are valid, whereas numerical models provide a more general description for complex or non-ideal datasets. Both analytical and numerical model types share the same result-writing code API. Consequently, fitted parameters, uncertainties, chi-squared values, AIC/BIC statistics, residuals and reconstructed curves can be stored consistently in the project. Automatic statistical model ranking using AIC/BIC is currently under development.

The exchange fitting backend also supports custom objective functions and staged fitting pipelines. Exchange models often contain strongly coupled parameters that result in poorly conditioned optimisation landscapes, particularly for multi-parameter or multi-condition fits. For complex models, the fitting procedure can therefore be decomposed into sequential minimisation steps in which selected parameters are varied while others are fixed, followed by refinement of the full or partial parameter set. This strategy improves convergence, maintains physically meaningful parameter bounds and allows each model to define an optimisation protocol appropriate to its assumptions.

A dedicated parameter-handling layer creates and initialises shared, context-local and observation-local parameters. For example, a CPMG model can fit shared exchange parameters such as $K_{ex}$ and exchange amplitude while assigning separate $R_{2,0}$ baselines to the data points recorded at different magnetic fields or experimental contexts. Similarly, numerical CEST models can fit global exchange parameters together with profile-specific variables, such as saturation time, B1 field strength, peak position, reference normalisation and relaxation rates. This separation of shared and local parameters allows exchange data acquired under different conditions to be fitted jointly without losing experiment-specific information.

The module also includes experiment-focused input selectors and preprocessing workflows. These workflows create analysis-ready data from raw spectra, generate intermediate tables, propagate errors and apply experiment-specific operations such as CEST profile normalisation, reference-point handling and automated dip picking. For CPMG and related dispersion experiments, intensity series can be converted to effective relaxation rates before model fitting. For CEST, complete intensity profiles can be fitted as array-valued observations using numerical two-state Bloch-McConnell propagation, with optional reference normalisation and B1 inhomogeneity handling.

Together, these features provide a flexible exchange-analysis framework that can support predefined, GUI-driven workflows while remaining extensible to new experiments and models. The separation between the Exchange Analysis Module and the Relaxation module allows AnalysisDynamics to accommodate the greater model diversity, input complexity and optimisation requirements of exchange experiments while preserving the same project-level data provenance, graphical integration and downstream compatibility.

#### Collections and DataTables

The standard series-analysis Collection hierarchy used for relaxation experiments was not sufficient for exchange experiments such as ZZ exchange, EXSY, CEST and DEST. In conventional relaxation analyses, including T1, T2 and hetNOE measurements, each assigned spin system typically corresponds to one observable that is followed across an experimental series (Supplementary Fig. 1a). CcpNmr assignment PIDs, such as A.A.25.H,N, identify the observed nuclei, and the data can be represented as an experiment-level Collection containing residue-level Collections, with peaks indexed by the series variable. By contrast, exchange experiments can generate multiple observables for the same spin system. In a two-state ZZ-exchange experiment[11], for example, the same residue can produce A→A, A→B, B→A and B→B peaks. These peaks share the same nuclei but differ in exchange meaning, so assignment alone is not sufficient to define the analysed quantity.

AnalysisDynamics addresses this issue by adding an observable-context level to the Collection hierarchy (Supplementary Fig. 1b). The assignment defines the observed spins, whereas the observable context defines the experiment-specific meaning of the peak. For ZZ-exchange or EXSY data, the context encodes a directed exchange pathway, such as A→B or B→A; for CEST or DEST data, it encodes a state-referenced or profile-specific observable. This avoids representing exchanging states as separate NMR chains, assignments or chemical-shift lists, which would duplicate spin-system identity and make exchange connectivity an indirect consequence of parallel assignments. Instead, all trajectories remain grouped within the same residue or spin-system context, while state or pathway information is encoded only where required.

To make these structures robustly identifiable within projects, AnalysisDynamics assigns registered namespaces to relevant Collection and DataTable objects (Supplementary Fig. 1c). A namespace records the application origin, semantic category and optional contextual tags associated with a stored object. The canonical namespace identifies the object family, such as an AnalysisDynamics exchange-series Collection or analysis DataTable, whereas tags provide non-identifying refinements such as experiment type, state label or exchange pathway. Namespaces are registered by native CcpNmr modules or plugins and can be resolved from stored objects when a project is reopened. This allows AnalysisDynamics to recognise, restore, filter and parse analysis objects without relying on user-defined names.

Together, observable contexts and namespaces extend the existing assignment model without changing the core CcpNmr data model. Observable contexts define the exchange meaning of individual peaks, while namespaces allow AnalysisDynamics to recognise analysis-specific Collections and DataTables and convert the corresponding stored data into structured input tables for relaxation or exchange analysis. The same infrastructure supports simple relaxation series,

ZZ-exchange/EXSY transfer pathways and CEST/DEST state-referenced profiles, while remaining extensible to assignment-light workflows, higher-state exchange models, plugin-defined analyses and future experiment types.

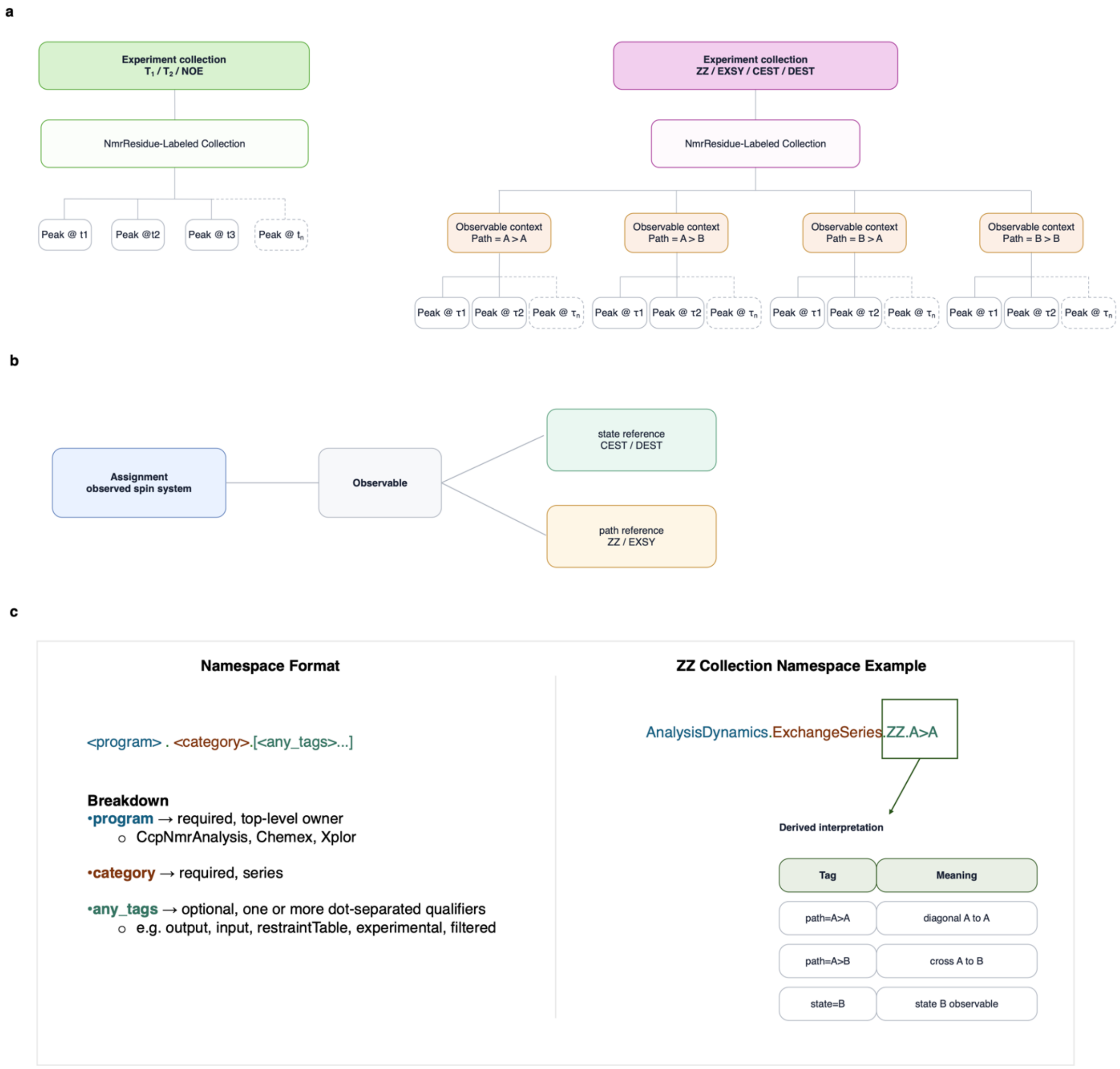

**Supplementary Figure 1. Collection hierarchy, observable contexts and CcpNmr DataTable namespaces in AnalysisDynamics. a**, Comparison of Collection object data hierarchy for standard relaxation and exchange experiments. In T1, T2 and hetNOE analyses, each assigned spin system is associated with a single observable that is followed across an experimental series. In exchange experiments such as ZZ exchange, EXSY, CEST and DEST, the same assigned spin system can give rise to multiple observables with distinct exchange meanings, such as state-specific or path-specific trajectories. **b**, Observable-context abstraction used to separate spin-system identity from experiment-specific interpretation. The assignment defines which nuclei are observed, whereas the observable context defines the meaning of the observation, including state references for CEST/DEST or directed transfer pathways for ZZ exchange and EXSY. **c**, Namespace scheme used to identify and store AnalysisDynamics Collection and DataTable object provenance.

**Supplementary Note 2. Plugin framework overview and design philosophy**

Within AnalysisDynamics, the new plugin system provides the mechanism by which native workflows and external analysis engines are exposed through a common project-level interface. The CcpNmr plugin framework is designed to support extensibility through a descriptor-driven and discovery-based architecture. Rather than enforcing inheritance from predefined base classes or requiring plugins to conform to rigid interfaces, the framework uses a declarative approach in which each plugin describes its entry points using a structured descriptor file. This design prioritises loose coupling between the core framework and plugin implementations, enabling external tools to be integrated without imposing constraints on their internal structure.

As a result, existing external NMR dynamics tools can be incorporated without requiring them to be rewritten as native CcpNmr modules (Fig. 3).

**Supplementary Note 3. Model Analysis (ModA)**

**Overview - architecture and implementation**

Model Analysis (ModA) computes $R_1$, $R_2$ and hetNOE values from a spectral-density description of molecular motion and estimates residue-specific dynamic parameters by numerical optimisation. The theoretical framework is standard and commonly referred to as the "Lipari-Szabo" or "ModelFree" approach[14,15].

The theoretical framework is implemented in a modular Python architecture that explicitly separates spectral-density theory, rotational diffusion models, relaxation-rate calculation and numerical optimisation[16]. This design enables clarity, extensibility and reproducibility, while maintaining computational efficiency for large datasets. Spectral-density models (Supplementary Table 1) are implemented as independent subclasses of an abstract base class, each defining a specific analytic expression for $J_\omega$ and an explicit list of optimised parameters[16]. These models are managed by a dedicated spectral-density handler, which acts as a registry and runtime selector for the active model set. This registry-based design allows new model parameterisations to be added without modifying the optimisation or rotational diffusion code sections. Rotational diffusion models (Supplementary Tables 2-4) are implemented as separate classes representing isotropic, axially symmetric and fully anisotropic tumbling. Each diffusion model encapsulates the conversion between diffusion constants and correlation times, as well as the calculation of orientation-dependent spectral-density weights. A diffusion-model handler orchestrates the execution of one or more diffusion models and supports automated model comparison using information-theoretic criteria.

Relaxation-rate calculations are handled by a central rate-management component that registers individual rate expressions (e.g., $R_1$, $R_2$, HETNOE, exchange and cross-correlation terms) under symbolic identifiers. Rate functions are implemented as stateless numerical routines and evaluated dynamically based on the spectral-density values produced by the active model. To ensure computational efficiency during iterative optimisation and Monte Carlo resampling, rate evaluations are cached using memoisation keyed by function identity and arguments, avoiding redundant calculations.

ModA also includes a dedicated parallel execution layer. Residue-wise datasets are partitioned into independent chunks and executed concurrently using Python's ProcessPoolExecutor, ensuring that each optimisation task runs in an isolated worker process. Completed results are collected asynchronously and merged into central data structures, enabling continuous progress monitoring and minimising overall runtime. This design allows the method to scale effectively with available CPU resources.

**High-level workflow**

A typical NMR dynamics analysis begins by assessing experimentally measured heteronuclear relaxation data, including $R_1$, $R_2$ and hetNOE values with associated uncertainties. From these data, an initial set of global parameters describing overall molecular tumbling is estimated, together with coarse residue-averaged measures of internal mobility. These global and local estimates are used solely to initialise the optimisation. Each residue is then analysed independently by fitting a hierarchy of spectral-density models that correspond to the standard model parameterisations (Supplementary Table 1). For each model, theoretical relaxation rates are calculated and the relevant dynamic parameters are optimised by minimising a goodness-of-fit objective function under physically motivated constraints. To improve robustness and to quantify parameter uncertainties, the optimisation is embedded within a Monte Carlo resampling procedure. The quality of each fit is assessed using the resulting $\chi^2$ values and derived model scores, enabling direct comparison between alternative spectral-density descriptions for a given residue. Finally, an optional global refinement step is performed in which the overall rotational correlation time is re-optimised using the full set of fitted residues to ensure internal consistency. The final output consists of residue-specific dynamic parameters, their uncertainties and the associated model selection metrics.

Steps:

1. Input experimental $R_1$, $R_2$ and hetNOE data with uncertainties.
2. Estimate initial global tumbling parameters and average motional amplitudes.
3. Compute initial residue-level estimates of internal motion.
4. For each residue and each spectral-density model:

   i) Calculate theoretical relaxation rates.
   ii) Optimise dynamic parameters under physical constraints.
   iii) Perform Monte Carlo resampling to estimate uncertainties.
   iv) Evaluate goodness of fit using $\chi^2$ and model scores.

5. Compare models and select the best description for each residue.
6. Optionally refine the global rotational correlation time using all residues.
7. Output final dynamic parameters with uncertainties.

#### Spectral-density functions

Heteronuclear NMR relaxation observables are interpreted using the Lipari–Szabo spectral-density formalism[17], in which the effects of molecular motion on relaxation are encoded through the frequency-dependent spectral-density function $J_\omega$. This approach separates overall rotational diffusion of the molecule from internal motions local to each bond vector, allowing internal dynamics to be characterised phenomenologically without assuming a detailed physical motional model. Within this framework, spectral-density functions are constructed as weighted sums of Lorentzian terms corresponding to the relevant frequency components of rotational diffusion, optionally supplemented by additional terms describing internal motion. A hierarchy of spectral-density parameterisations is implemented, corresponding to the standard Lipari–Szabo models commonly denoted models 1–4 (Supplementary Table 1). These models differ in the number and nature of internal motion parameters, including single or multiple order parameters and effective internal correlation times, with optional additive chemical-exchange contributions to the transverse relaxation rate.

The code for spectral-density calculations is kept independent of the relaxation-rate equations, allowing the same spectral-density models to be combined consistently with different diffusion descriptions and relaxation observables.

| Model Name | Spectral Density Function | Optimised Parameter |
|---|---|---|
| 1 | $J(w) = \frac{2}{5}\left[S^2 \sum_{i=1}^{n} \frac{c_i \cdot \tau_i}{1 + (\tau_i \omega)^2}\right]$ | $\tau_i,\ S^2$ |
| 2 | $J(w) = \frac{2}{5}\left[S^2 \sum_{i=1}^{n} \frac{c_i \cdot \tau_i}{1 + (\tau_i \omega)^2} + \frac{(1 - S^2)\tau}{1 + (\tau\omega)^2}\right]$ | $\tau_i,\ S^2,\ \tau_e$ |
| 3 | $J(w) = \frac{2}{5}\left[S^2 \sum_{i=1}^{n} \frac{c_i \cdot \tau_i}{1 + (\tau_i \omega)^2}\right], R_{ex}$ | $\tau_i,\ S^2,\ R_{ex}$ |
| 4 | $J(w) = \frac{2}{5}\left[S^2 \sum_{i=1}^{n} \frac{c_i \cdot \tau_i}{1 + (\tau_i \omega)^2} + \frac{(1 - S^2)\tau}{1 + (\tau\omega)^2}\right], R_{ex}$ | $\tau_i,\ S^2,\ \tau_e,\ R_{ex}$ |

Supplementary Table 1. Spectral-density models and associated parameterisations implemented in ModA

#### Rotational diffusion models

Overall molecular tumbling is described using rotational diffusion models of increasing complexity. Three diffusion models are implemented: isotropic, axially symmetric and fully anisotropic diffusion[17]. In the isotropic model, global tumbling is characterised by a single rotational correlation time and does not require structural information (Supplementary Table 2). The axially symmetric model introduces two diffusion constants corresponding to motion parallel and perpendicular to a unique molecular axis and requires knowledge of the orientation of each N–H bond vector relative to the diffusion tensor

(Supplementary Table 3). The fully anisotropic model generalises this description to three independent diffusion constants, yielding five correlation times and fully orientation-dependent spectral-density weights (Supplementary Table 4).
For each diffusion model, analytic relationships are used to convert diffusion constants into the corresponding set of rotational correlation times and weighting coefficients. Initial estimates of the diffusion parameters are generated from a global tumbling time and refined during optimisation. This hierarchical treatment allows systematic evaluation of isotropic and anisotropic descriptions of molecular tumbling within a unified framework.

For a given residue, diffusion model and spectral-density parameterisation, theoretical relaxation rates are generated using a forward calculation that maps model parameters to predicted observables. Model parameters define the spectral-density function $J_\omega$ via the selected spectral-density model. For each spectrometer field present in the dataset, $J_\omega$ is evaluated at the discrete angular frequencies required by heteronuclear relaxation theory, including zero frequency and the relevant proton and heteronuclear single-frequency and combination-frequency terms.
All field-dependent physical quantities, such as Larmor frequencies and relaxation prefactors arising from dipolar and chemical shift anisotropy (CSA) interactions, are precomputed once per field and treated as fixed inputs. Consequently, per-residue optimisation of the CSA tensor, as proposed in Fushman et al. 1999[18], is currently not implemented. The resulting spectral-density values are passed to the appropriate relaxation-rate expressions to compute theoretical longitudinal relaxation rates, transverse relaxation rates and hetNOE values. When a spectral-density model includes chemical exchange, the exchange contribution is added explicitly to the transverse relaxation rate without modifying the spectral-density function itself. Predicted relaxation rates are concatenated across all available fields and observables to form a model prediction vector, which is supplied to the objective function during numerical optimisation.

Isotropic rotational diffusion

| $\boldsymbol{i}$ | $\boldsymbol{i = 1}$ |
|---|---|
| $\boldsymbol{c_i}$ | $C_1 = 1$ |
| $\boldsymbol{\tau_i}$ | $\tau_1 = \frac{1}{6R}$ |

Supplementary Table 2. Isotropic diffusion terms used in the ModA spectral-density expressions

With:
$R = \frac{kT}{8\pi\eta r^3}$

Axially Symmetric rotational diffusion

| $i$ | $i = 1$ | $i = 2$ | $i = 3$ |
|---|---|---|---|
| $c_i$ | $C_1 = \frac{1}{4}(3x^2 - 1)^2$ or $\frac{1}{4}(3\cos^2\theta - 1)^2$ | $C_2 = 3x^2(1 - x^2)$ or $3\sin^2\theta\cos^2\theta$ | $C_3 = \frac{3}{4}(x^2 - 1)^2$ or $\frac{3}{4}\sin^4\theta$ |
| $\tau_i$ | $\tau_1^{-1} = 6D_\perp$ | $\tau_2^{-1} = D_\perp + 5D_\parallel$ | $\tau_3^{-1} = 4D_\perp + 2D_\parallel$ |

Supplementary Table 3. Axially symmetric terms used in the ModA spectral-density expressions

With $x = \cos\theta$

where $\theta$ is the angle between the N-H vector and the unique axis of the gyroscopic diffusion tensor.

Anisotropic rotational diffusion

| $i$ | $i = 1$ | $i = 2$ | $i = 3$ | $i = 4$ | $i = 5$ |
|---|---|---|---|---|---|
| $c_i$ | $C_1 = 6y^2z^2$ | $C_2 = 6x^2z^2$ | $C_3 = 6x^2y^2$ | $C_+ = d + e$ | $C_- = d - e$ |
| $\tau_i$ | $\tau_1^{-1} = 4D_{xx} + D_{yy} + D_{zz}$ | $\tau_2^{-1} = D_{xx} + 4D_{yy} + D_{zz}$ | $\tau_3^{-1} = D_{xx} + D_{yy} + 4D_{zz}$ | $\tau_+^{-1} = 6[R + (R^2 - L^2)^{\frac{1}{2}}]$ | $\tau_-^{-1} = 6[R - (R^2 - L^2)^{\frac{1}{2}}]$ |

Supplementary Table 4. Anisotropic terms used in the ModA spectral-density expressions

with:

$$d = \frac{1}{2}[3(x^4 + y^4 + z^4) - 1]$$

$$e = \frac{1}{6}[\delta_1(3x^4 + 6y^2z^2 - 1) + (3y^4 + 6x^2z^2 - 1) + (3z^4 + 6x^2y^2 - 1)]$$

$$\delta_1 = \frac{(D_{xx} - R)}{(D_{yy} - L^2)^{\frac{1}{2}}} \qquad \delta_2 = \frac{(D_{yy} - R)}{(D_{yy} - L^2)^{\frac{1}{2}}} \qquad \delta_3 = \frac{(D_{zz} - R)}{(D_{yy} - L^2)^{\frac{1}{2}}}$$

$$R = \frac{1}{3}(D_{xx} + D_{yy} + D_{zz})$$

$$L^2 = \frac{1}{3}(D_{xx}D_{yy} + D_{xx}D_{zz} + D_{yy}D_{zz})$$

### Differential evolution optimisation

Differential Evolution (DE), implemented through LMFIT and SciPy optimisation routines, forms the core of the ModA global optimisation strategy[3,5]. Conceptually, DE operates by allowing a population of trial solutions to “explore” the parameter space collectively. Each generation of solutions is perturbed by adding scaled differences between randomly chosen population members, and these proposals are blended with existing candidates before being evaluated against the objective function. Trial solutions that offer improvement are retained, and those that do not are discarded. This simple cycle of mutation, crossover and selection enables the population to adaptively refine its search through the often-complex landscapes encountered in nonlinear model fitting. Importantly, DE requires no gradients, no assumptions of smoothness and no tuning beyond a few intuitive control parameters, making it especially suited for problems where the structure of the optimisation surface is not known a priori.

We use Differential Evolution (DE) as the global optimisation strategy in ModA because it provides a population-based, gradient-free search method suitable for bounded nonlinear optimisation problems. This is useful for Lipari-Szabo analysis, where order parameters, internal correlation times and diffusion-related parameters can be correlated and where the objective function may depend on parameter bounds and starting conditions. Compared with exhaustive grid searches, DE avoids the rapid scaling of computational cost with parameter number, while providing broader exploration than purely local minimisation. In ModA, DE is therefore used to identify plausible regions of parameter space before subsequent refinement and model assessment.

In practice, Differential Evolution exposes a large number of algorithmic controls, such as population size, mutation and recombination rates, convergence tolerances and choice of mutation strategy, that influence the balance between global exploration and local refinement. To make these capabilities accessible while simultaneously ensuring robust performance across diverse relaxation datasets, we provide a set of curated parameter settings (pre-sets) that bundle empirically optimised parameter values for different analysis goals. A low-accuracy pre-set enables rapid screening or exploratory model assessment, using modest population sizes and relaxed tolerance thresholds to characterise the broad structure of the parameter landscape at minimal computational cost. The medium- and high-accuracy pre-sets progressively tighten convergence criteria, increase evolutionary depth and employ more exploitative strategies (e.g., rand-to-best variants) suited to the subtle curvature of Lipari–Szabo $\chi^2$ surfaces. For advanced users, a custom mode exposes the full DE configuration, including strategy selection, population initialisation schemes (Sobol, Halton, Latin hypercube), mutation dithering ranges, population scaling and polishing behaviour, allowing fine-grained control of the optimisation process when tackling particularly challenging parameter regimes. This tiered design provides a practical workflow: rapid global exploration when first approaching a dataset, followed by progressively more stringent refinement once a promising region of parameter space has been identified. By structuring these choices as well-defined pre-sets, we preserve the flexibility inherent to DE while ensuring reproducible, user-friendly optimisation behaviour appropriate for Lipari–Szabo analysis.

#### Chi-Squared $\chi^2$

The objective function used throughout this work differs intentionally from the conventional least-squares $\chi^2$ often employed in relaxation fitting. Standard $\chi^2$ assumes a uniform additive-Gaussian noise model with accurately known uncertainties, but actual experimental $R_1$, $R_2$ and hetNOE data rarely satisfy these conditions simultaneously. Each observable carries its own noise structure: $R_1$ and $R_2$ values tend to exhibit predominantly additive uncertainty, whereas HETNOE intensities are well-known to follow multiplicative (log-normal–like) behaviour, with errors that scale with signal magnitude. Moreover, reported experimental uncertainties can vary widely in reliability; underestimated $\sigma$ values produce artificially inflated $\chi^2$, whereas overly conservative σ values lead to overly flat objective landscapes, masking genuine differences between models. These effects can destabilise optimisation, distort model selection and degrade uncertainty estimates if left unaddressed.

In Lipari-Szabo spectral-density analysis, experimental uncertainties directly determine the weighting of each relaxation observable in the objective function. Unrealistically small σ values, for example from overconfident exponential fits or anomalously high signal-to-noise estimates, can cause individual data points to dominate the fit. Conversely, excessively large σ values flatten the contribution of an observable to the objective function, effectively removing that measurement from influencing the fit. In LS analysis this often manifests as poorly constrained $S^2$, overly permissive $\tau_e$ values, or an inability to discriminate between competing motional models (Supplementary Table 1). For heteronuclear NOE, which normally functions the strongest discriminator between fast internal motion and slower global tumbling, inflated uncertainties can cause the optimiser to drift into degenerate regions of parameter space where multiple models yield comparable $\chi^2$. This leads to unreliable model selection, inflated parameter uncertainties and a loss of sensitivity to genuine dynamical features such as sub-nanosecond internal flexibility or weak anisotropy in the rotational diffusion tensor. Together, these effects mean that naïvely using the reported uncertainties, whether underestimated or overestimated, can distort both the numerical stability and the physical fidelity of Lipari–Szabo analysis. The noise-adaptive $\chi^2$ introduced here is specifically designed to mitigate these undesired effects and restore a well-conditioned optimisation landscape.

To account for these realities of the Lipari-Szabo modelling, we implemented a group-aware, noise-adaptive $\chi^2$ approach in ModA. Each observable type, i.e. $R_1$, $R_2$, HETNOE, is treated as a separate statistical group. For each group the algorithm automatically selects either an additive Gaussian model or a log-Gaussian model, depending on whether residual magnitude correlates with prediction magnitude. This allows the objective function to reflect the empirical noise behaviour of the data rather than imposing a one-size-fits-all model. Within each group, nominal errors are stabilised using adaptive absolute- and relative floors, preventing tiny or inflated uncertainties from dominating the fit. For robustness against occasional experimental outliers, we optionally apply a Huber-loss function[19] to the standardised residuals, preserving quadratic behaviour for well-behaved points while linearly down-weighting extreme deviations. The resulting $\chi^2$ is still additive across data points and groups, but its effective weighting dynamically reflects both the noise structure and the reliability of uncertainties within each observable class (Supplementary Fig. 2).

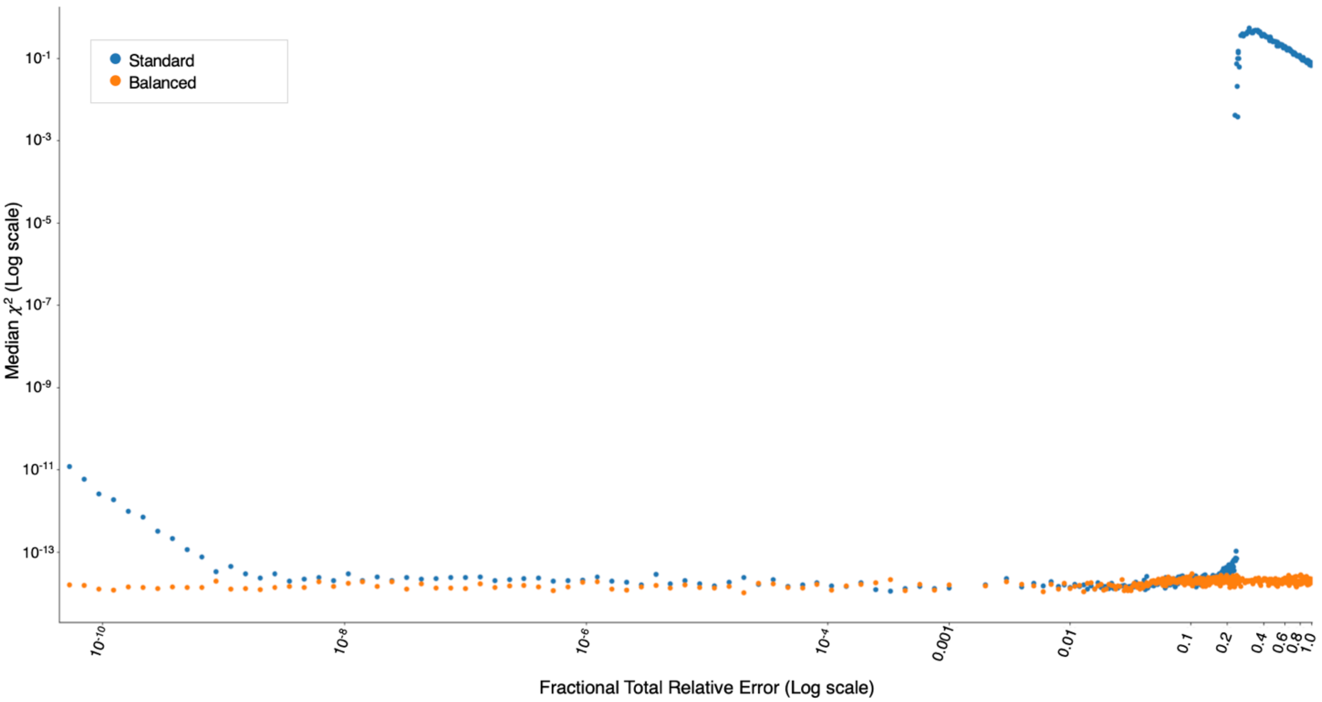

**Supplementary Figure 2. Uncertainty scaling and χ² stability analysis**. The plot compares standard calculations (blue data points) and balanced χ² calculations (orange data points) across the same uncertainty range, showing that standard χ² values are highly sensitive to extreme uncertainty scaling, whereas the balanced calculation remains comparatively stable over the experimentally relevant regime.

### Error estimation: Monte Carlo, Latin hypercube sampling

To quantify parameter uncertainties and assess the stability of the Lipari–Szabo fits, we employ a Monte Carlo resampling approach in which synthetic datasets are generated by perturbing the experimental relaxation rates within their measured uncertainties. Rather than drawing noise independently for each data point using simple Gaussian deviates, which can lead to uneven sampling and poor coverage in high-dimensional settings, we generate the perturbations using a Latin Hypercube Sampling (LHS) scheme[20]. LHS stratifies the unit hypercube into equal-probability segments and draws one sample from each, ensuring that each Monte Carlo replicate explores a distinct and well-distributed region of the underlying uncertainty space. This substantially improves the efficiency of the uncertainty estimation, particularly for heterogeneous datasets, such as those involving $R_1$, $R_2$ and hetNOE rates with different noise properties.

Each pseudo-dataset is constructed by sampling from a truncated normal distribution centred on the experimental value and bounded symmetrically at ±5σ, optionally enforcing physical constraints such as rate positivity. Importantly, we retain the original reported σ values for weighting in the χ² calculation so that each synthetic dataset reflects only the uncertainty in the measured rates, not an altered noise model. The first minimisation is performed on the unperturbed data, and subsequent Monte Carlo iterations begin from slightly perturbed parameters, allowing the optimiser to re-explore nearby basins of attraction without drifting into extreme regions. Re-running the full fit on each pseudo-dataset yields a collection of parameter vectors whose variability directly reflects the sensitivity of the model to measurement noise. Final parameter uncertainties are then estimated using robust statistics, such as the median absolute deviation, which are less susceptible to occasional unstable minimisations or non-Gaussian parameter distributions.

This combination of truncated-noise Monte Carlo, Latin hypercube stratification and full-refit propagation produces uncertainty estimates that are both statistically principled and numerically stable. It respects the heterogeneous noise structure of relaxation observables, avoids degeneracies arising from poor random sampling and captures correlations between fitted parameters that cannot be recovered from linearised error propagation. In the context of Lipari–Szabo analysis where parameters may be strongly coupled and the optimisation landscape can exhibit multiple basins, this approach provides a more reliable characterisation of parameter precision and robustness than analytic approximations or naïve bootstrapping.

**Model selection and confidence scores**

To identify the most appropriate Lipari–Szabo model for each residue, we evaluate all candidate models using a suite of likelihood-based information criteria. Because each model is fitted by minimising our $\chi^2$-like objective, the Akaike Information Criterion (AIC) and Bayesian Information Criterion (BIC) can be expressed in their reduced forms as simple penalised sums of $\chi^2$, with additional penalties that depend on the number of free parameters k and the number of observations n. For a model with chi-square value $\chi^2$, these criteria take the form

$$AIC = x^2 + 2k \tag{1}$$

$$BIC = x^2 + kln\,(n) \tag{2}$$

which naturally balance model fit against model complexity. Because the number of experimental datapoints per residue is often modest, we also compute the small-sample corrections AICc and BICc[21], which include an additional term that penalises over-parameterised models when n - k - 1 is small. These corrected forms reduce the risk of favouring higher-order Lipari–Szabo models simply due to their additional degrees of freedom. For each residue, the model with the lowest information criterion value is taken as the best-supported model for that dataset.

However, information criteria computed on a single fit can underestimate the uncertainty in model selection when measurement noise is substantial or when two models produce comparably good fits. To capture this variability, we extend model selection into a Monte Carlo confidence framework. For each residue, we generate many pseudo-datasets by perturbing the experimental relaxation rates according to their reported uncertainties using a truncated normal distribution. Each replicate dataset is then refitted across all candidate models, and the model scores, i.e., AIC, BIC, or whichever criterion is chosen by the user, are recalculated. For each iteration, the model with the lowest score is recorded. Aggregating these results across all Monte Carlo replicates yields a confidence score for each model, defined as the fraction of replicates in which that model is selected as the best. This yields a probability-like measure of model support, analogous to bootstrap model frequencies used in statistical inference.

This dual strategy of using penalised likelihood for objective model comparison and Monte Carlo frequencies for confidence estimation provides a far more nuanced view of model selection compared to using AIC/BIC alone. Models that are decisively superior across noise realisations receive high confidence scores, whereas residues with ambiguous dynamics or limited data naturally exhibit more diffuse model distributions. This enables direct identification of both well-determined and uncertain residues in terms of dynamical model determination and supports downstream error propagation in parameters and derived timescales. The result is a model selection framework that is statistically grounded, noise-aware and robust to the complexities of real NMR relaxation datasets. To facilitate interpretation, we provide a visual representation of the model-selection results using intuitive colour gradients, together with a complete dataset of per-residue scores, confidence values and fitted parameters for full transparency and downstream analysis.

**Parameter initialisation: Global Correlation Time $\tau_c$**

An initial estimate of the overall rotational correlation time, $\tau_c$, is obtained prior to residue-level optimisation in order to provide physically reasonable starting values for the diffusion parameters. $\tau_c$ is estimated directly from experimental relaxation data using a modular estimation framework that operates on grouped $R_1$ and $R_2$ measurements at each spectrometer field strength. The default implementation follows the approach of Fushman and co-workers[22], in which $\tau_c$ is derived from the ratio of longitudinal and transverse relaxation times under the assumption that internal motions are limited and that overall tumbling dominates the relaxation behaviour.

To reduce bias from residues exhibiting atypical dynamics, the $\tau_c$ estimation incorporates optional data-filtering steps applied independently at each field. Residues with low hetNOE values are excluded using a user-defined HETNOE threshold, thereby removing sites likely affected by substantial internal flexibility. In addition, a rate-relative-deviation filter is applied to identify and exclude outliers based on the difference between normalised deviations of $R_1$ and $R_2$ from their global central values. This second filter suppresses contributions from residues with anomalous relaxation behaviour that could distort the global $\tau_c$ estimate.

Following filtering, per-residue $\tau_c$ values are computed from the remaining $R_1$ and $R_2$ data at each field using an analytic expression relating the ratio of relaxation times to the heteronuclear Larmor frequency. The median is then used to obtain a field-specific $\tau_c$ value, and the final initial $\tau_c$ is taken as the average of these estimates across all available spectrometer fields. The resulting $\tau_c$ serves exclusively as an initial global parameter for subsequent optimisation and is not interpreted as a final dynamic result; a refined global $\tau_c$ is obtained later through joint optimisation against the full set of fitted relaxation data.

Using the same filtered dataset, an initial estimate of the average generalised order parameter $\langle S^2 \rangle$ is computed from the product $R_1 \cdot R_2$ following Kneller et al[23]. A robust central estimator is applied to the distribution of $R_1$ $R_2$ values, using either a trimmed mean or the median depending on the selected method. The resulting central value is normalised by the maximum observed $R_1 \cdot R_2$ product; the square root of this value then yields $\langle S^2 \rangle$. This estimate provides a global measure of motional restriction and is used solely to initialise residue-level order parameters prior to full spectral-density–based fitting.

**Parameter initialisation: Effective Correlation Time $\tau_e$**

Accurate initialisation of the effective internal correlation time $\tau_e$ is essential for stable Lipari–Szabo optimisation, as $\tau_e$ is strongly coupled to both the order parameter $S^2$ and the global tumbling correlation time $\tau_c$. Poor seeds can produce unrealistic search ranges, slow convergence, or spurious minima—particularly when residues differ substantially in dynamical character. To address this, the framework provides two complementary strategies for defining initial $\tau_e$ values and optimisation bounds: a global $\tau_e$ initialisation and an adaptive per-residue $\tau_e$ initialisation.

In the global initialisation approach, $\tau_e$ is derived solely from the overall tumbling time $\tau_c$ using predefined fold factors, f. The initial value is set as $\tau_e = \tau_c/f$, with lower and upper bounds similarly generated as scaled fractions of $\tau_c$. This yields a physically reasonable but uniform $\tau_e$ window applied identically to all residues. The method is simple, computationally light and often sufficient for exploratory analyses. However, because it ignores residue-specific motional amplitudes, the resulting bounds can be overly broad for rigid sites, inflating search space, or too narrow for flexible sites, thus biasing the optimisation away from slow internal motions.

In our adaptive per-residue approach, $\tau_e$ bounds are computed for each residue from its fitted order parameter $S^2$ and the global $\tau_c$, ensuring that rigid residues receive narrow $\tau_e$ windows while flexible residues are allowed broader, slower-timescale intervals. The initial $\tau_e$ value is placed geometrically between these bounds, preserving the multiplicative scaling inherent to internal motions and preventing bias towards extremes of the parameter space.

To define the admissible upper limit, we impose a ceiling that contracts smoothly as $S^2$ approaches unity (rigidity) while expanding for more flexible residues. This is expressed as:

$$\tau_e, max = min(\tau_{cap}, c \cdot \tau_c \cdot (1 - S^2)^{\beta}) \quad (3)$$

where the factor $c \cdot \tau_c$ sets the residue-independent upper scale and the term $(1 - S^2)^{\beta}$ modulates this range in a physically interpretable manner: small internal amplitudes ($S^2 \approx 1$) suppress $\tau_{e,max}$, whereas larger amplitudes permit slower correlation times. The exponent β determines how sensitively $\tau_{e,max}$, responds to $S^2$, allowing smooth tuning of the narrowing behaviour.

To further refine these per-residue windows, we introduce a data-driven bias parameter derived from percentile-based rankings of $R_2/R_1$ ratios across the data obtained across magnetic fields, if available, optionally incorporating HETNOE information and simple exchange flags. Because $R_2/R_1$ increases systematically with motional restriction, these rankings provide a robust empirical proxy for residue-level rigidity. The ranked rigidity index $p_i$ is then mapped into a bounded interval [$w_{min}$, $w_{max}$] to produce a smooth, residue-specific bias that shifts $\tau_e$ initialisation toward either the fast or slow end of the search window:

$$w_i = wmin + (wmax - wmin) \cdot p_i. \quad (4)$$

This linear mapping ensures that residues exhibiting high $R_2/R_1$ (restricted motion) preferentially initialise $\tau_e$ near the lower bound, whereas dynamically flexible residues initialise closer to the upper bound. Because $w_i$ parameterises a geometric interpolation between $\tau_{e,min}$ and $\tau_{e,max}$, it preserves the log-scaled structure of internal timescales and avoids distortions that would arise from linear interpolation.

Together, these adaptive elements provide a residue-specific alternative to global $\tau_e$ initialisation schemes, which impose a single set of bounds and starting conditions across all residues. Global schemes necessarily under-constrain rigid sites and over-constrain flexible ones, promoting convergence to artefactual minima and increasing sensitivity to noise, especially under Differential Evolution and other global optimisers. In contrast, residue-specific $\tau_e$ adaptation produces physically coherent search spaces that capture the inherent heterogeneity of internal motions, leading to markedly improved optimisation stability, reduced parameter degeneracy and more reliable Lipari–Szabo model fits across the entire protein.

**Parameter finalisation: Global Correlation Time $\tau_c$**

After residue-level Lipari-Szabo optimisation, we perform a final global refinement of the overall tumbling description to improve internal consistency across fitted residues. This step recognises that $\tau_c$ is a global property of the molecule, whereas per-residue fits may introduce small numerical deviations due to noise, model heterogeneity or local parameter correlations. The refinement uses a single multiplicative diffusion-scaling factor, $s$, applied uniformly to the fitted rotational diffusion eigenvalues $(D_1, D_2, D_3)$ for all residues.

To keep the scaling factor positive during optimisation, ModA does not fit $s$ directly. Instead, it fits the unconstrained variable $\alpha$ , defined as $\alpha = \log s$

$$s = \exp(\alpha) \tag{5}$$

Bounds are specified as a narrow fractional range around $s = 1$, typically $0.9 \leq s \leq 1.1$ and are converted to corresponding bounds on corresponding bounds on $\alpha$. This prevents large corrections while allowing small systematic deviations from the residue-level fits to be reconciled. For each residue *i*, the base diffusion tensor $D_{\mathrm{i}}^{(0)}$ obtained from its individual optimisation is rescaled as

$$D_{\mathrm{i}}(s) = s \cdot D_{\mathrm{i}}^{(0)} \tag{6}$$

after which the corresponding rotational timescales and geometric tensors are recomputed. All internal Lipari–Szabo parameters remain fixed during this step, ensuring that the refinement adjusts only the global tumbling behaviour without perturbing fitted internal dynamics.

A global least-squares objective is then assembled from the standardised residuals between the experimental relaxation observables, $R_1$, $R_2$ and hetNOE and the corresponding values back-calculated after diffusion scaling. The optimal scaling factor is obtained by minimising

$$s_{\mathrm{opt}} = \arg \min_{s} \sum_{i} \sum_{j} \left(\frac{R_{i,j} - \hat{R}_{i,j}(s)}{\sigma_{i,j}}\right)^2 \qquad (7)$$

where *i* indexes residues, *j* indexes relaxation observables and $\hat{R}_{i,j}(s)$ denotes the back-calculated value obtained using the scaled diffusion components $D_i(s)$. Following optimisation, per-residue effective $\boldsymbol{\tau}_c$ values are recomputed from the scaled diffusion components and the final global $\boldsymbol{\tau}_c$ is reported as their median. This provides a robust summary of the globally scaled tumbling description while reducing artefactual variability introduced during residue-wise fitting.

**Tensor handling and bond-vector geometry**

For diffusion models requiring structural information, atomic coordinates are obtained from standard Protein Data Bank (PDB) or mmCIF files and processed through a unified structural interface. Structures are validated to ensure the presence of backbone nitrogen–hydrogen vectors suitable for relaxation analysis and a single structural model is selected when ensemble coordinates are provided.

Overall molecular shape can be characterised using the radius-of-gyration tensor G[24], defined as the second moment of the atomic coordinate distribution relative to the centre of mass,

$$\mathbf{G} = \langle (\mathbf{r}_i - \mathbf{r}_{\mathrm{cm}})(\mathbf{r}_i - \mathbf{r}_{\mathrm{cm}})^{\mathrm{T}} \rangle_i \qquad (8)$$

where $\mathbf{r}_i$ are atomic coordinates and $\mathbf{r}_{\mathrm{cm}}$ is the molecular centre of mass. Eigenvalue decomposition of G yields three orthonormal eigenvectors and associated eigenvalues $\lambda_1$, $\lambda_2$, $\lambda_3$, which provide a compact description of molecular anisotropy. The relative magnitudes of these eigenvalues are used to classify the molecule as isotropic, axially symmetric or fully anisotropic, guiding selection of the appropriate rotational diffusion model.

For axially symmetric systems, the principal diffusion axis is identified as the eigenvector associated with the largest eigenvalue. Effective geometric estimates of the parallel and perpendicular diffusion components are derived as

$$R_{\parallel} = \lambda_{max}, \qquad R_{\perp} = \frac{1}{2}(\lambda_2 + \lambda_3) \qquad (9)$$

with corresponding initial diffusion coefficients defined as $D_{\parallel} \propto R_{\parallel}^{-1}$ and $D_{\perp} \propto R_{\perp}^{-1}$. These quantities are used exclusively as physically motivated initial estimates and do not constitute explicit hydrodynamic modelling.

Backbone N–H bond vectors $\mathbf{u}_i$ are constructed from nitrogen and amide hydrogen atomic coordinates. In axially symmetric diffusion models, the orientation of each bond vector relative to the principal diffusion axis $\mathbf{e}_\parallel$ is given by

$$\cos\,\theta_i = \hat{\mathbf{u}}_i \cdot \hat{\mathbf{e}}_\parallel \tag{10}$$

where hats denote normalised vectors. For fully anisotropic diffusion models, bond-vector orientations are represented by their direction cosines with respect to the diffusion-frame eigenvectors,

$$(\ell_i, m_i, n_i) = \mathbf{E}^{\mathrm{T}}\mathbf{u}_i, \tag{11}$$

where E is the matrix of eigenvectors of the gyration tensor. Structural information is used exclusively to define rotational diffusion geometry and bond-vector orientation and does not impose additional assumptions on internal dynamics.

### Supplementary Note 4. Consistency Test Metrics

#### Introduction

Relaxation data acquired at multiple magnetic field strengths must be internally consistent before being combined in quantitative analyses[25]. While multi-field measurements enhance sensitivity to molecular dynamics and improve parameter identifiability, systematic differences arising from sample conditions or acquisition protocols can introduce field-dependent biases. Consistency testing mitigates this risk by comparing field-independent quantities derived from the relaxation rates across magnetic fields, thereby identifying datasets that may compromise joint model fitting. This procedure provides an essential quality control step prior to detailed dynamical analysis.

#### J(0)–J(0) field consistency test

Consistency testing is performed using the zero-frequency spectral density J(0), estimated independently at each magnetic field using the reduced spectral density formalism. J(0) captures low-frequency motional and exchange contributions that predominantly affect $R_2$ and, to first order, should be independent of magnetic field strength. For each residue, J(0) values computed at different fields are therefore expected to coincide within experimental uncertainty. Pairwise J(0)–J(0) scatter plots provide a direct visual assessment of consistency, where agreement is indicated by points lying close to the identity line. Systematic deviations from linearity or clusters of outliers reveal field-dependent biases that would compromise joint fitting of multi-field relaxation data.

#### Q-Q, PCA and Q-scores

Quantile–quantile (Q–Q) analysis

To complement residue-wise comparisons, J(0) distributions obtained at different fields are further examined using quantile–quantile (Q–Q) plots. In this representation, corresponding quantiles of the two J(0) distributions are plotted against one another. If the datasets are statistically consistent, the quantiles follow the identity line across the full dynamic range. Deviations from linearity indicate differences in distribution shape, such as systematic shifts, changes in spread, or the presence of field-specific outliers. Q–Q analysis therefore provides a global, distribution-level assessment of consistency that is less sensitive to individual residues than direct scatter plots.

PCA analysis

While pairwise J(0)–J(0) comparisons are sufficient for two magnetic fields, consistency assessment becomes less transparent as additional fields are included. To address this, J(0) values from all fields are analysed jointly using principal component analysis (PCA) after standardisation. PCA identifies the dominant modes of variation across fields and residues, enabling inconsistencies to be detected in a reduced-dimensional representation. In the absence of systematic field-dependent effects, the majority of variance is captured by a single principal component, with residues clustering near the origin in higher-order components.

Residues that deviate from the main PCA cluster indicate field-dependent behaviour inconsistent with a common J(0) value. To quantify such deviations, a per-residue Q-score is computed from the reconstruction error obtained by projecting the standardised J(0) data onto the full PCA model and transforming back to the original space. Large Q-scores correspond to residues that are poorly described by the dominant variance structure and are therefore likely to reflect experimental inconsistencies or anomalous relaxation behaviour. Plotting Q-scores as a function of residue index enables rapid identification of outliers, which may be excluded or examined further prior to dynamical modelling.

Several limitations of the present consistency analysis should be noted. First, J(0) estimates inherit uncertainties from $R_1$, $R_2$ and HETNOE measurements and are therefore sensitive to experimental noise, particularly for residues with weak NOEs or poorly determined $R_2$ values. Second, because all tests probe low-frequency contributions to relaxation, they cannot distinguish between true dynamic heterogeneity and field-dependent experimental artefacts. Third, PCA-based metrics identify statistical outliers but do not by themselves provide a physical explanation for inconsistency. Consequently, residues flagged by J(0)-based tests should be interpreted conservatively and examined in the context of experimental conditions, structural environment and relaxation fitting results.

**Benchmark on real BMRB Datasets**

We developed an automated analysis workflow to evaluate the internal consistency of multi-field $^{15}N$ relaxation datasets remediated from the Biological Magnetic Resonance Data Bank (BMRB). For each BMRB entry, zero-frequency spectral density values J(0) were computed independently for each reported spectrometer frequency and compared using a combination of pairwise distributional statistics and multivariate analysis. Consistency across fields within each entry was quantified using Kolmogorov–Smirnov statistics, quantile–quantile agreement metrics, fractional-change dispersion and principal component analysis performed separately for each dataset. These measures were normalised and combined into a single dataset-level score, allowing BMRB entries to be ranked and classified according to their internal consistency.

**Consistency test multi-level scoring**

For each dataset, a zero-frequency spectral density measure, J(0), is estimated independently at each magnetic field and used as a field-independent reference quantity for assessing internal consistency across measurements. Comparisons are then performed between the resulting field-specific J(0) distributions to identify systematic deviations indicative of field-dependent biases.

Kolmogorov–Smirnov statistic

The two-sample Kolmogorov–Smirnov (KS) statistic is computed between the two J(0) vectors:

$$\mathrm{KS}(f_i, f_j) = \sup_x \left| F_{f_i}(x) - F_{f_j}(x) \right| \tag{12}$$

where $F_{f_i}(x)$and $F_{f_j}(x)$ are the empirical cumulative distribution functions of the two distributions.

Quantile–quantile comparison and Q–Q $R^2$

For each frequency pair, matched empirical quantiles are computed:

$$Q_f(p_k) = \mathrm{quantile}(\mathbf{J}_f, p_k), p_k \in [0,1] \tag{13}$$

The deviation from perfect agreement is quantified using an $R^2$-like statistic:

$$R^2_{\mathrm{QQ}} = 1 - \frac{\sum_k [Q_{f_y}(p_k) - Q_{f_x}(p_k)]^2}{\sum_k [Q_{f_y}(p_k) - \overline{Q}_{f_y}]^2} \tag{14}$$

where

$$\overline{Q}_{f_y} = \frac{1}{K} \sum_k Q_{f_y}(p_k)$$

This score is computed for each frequency pair, treating fj as the reference quantile distribution, and is then summarised at the dataset level. Because the statistic is $R^2$-like rather than a bounded correlation coefficient, strongly inconsistent frequency pairs can yield negative values.

Fractional change and MADQ statistic

Residue-wise fractional changes between two frequencies are computed as:

$$\Delta_{ij}(n) = \frac{|J_{f_i}(0)_n - J_{f_j}(0)_n|}{|J_{f_i}(0)_n|} \tag{15}$$

The distribution of fractional changes is summarised using the median absolute deviation of quantiles (MADQ ). Let $\tilde{\Delta}$ denote the median of $\Delta_{ij}$ and let $Q_\Delta(p_k)$ denote its quantiles. Then:

$$\mathrm{MAD}_Q = \mathrm{median}_k \left| Q_\Delta(p_k) - \tilde{\Delta} \right| \tag{16}$$

PCA

For each dataset, a matrix of residue-wise J(0) values is assembled:

$$\mathbf{X} \in \mathbb{R}^{N \times F} \tag{17}$$

where N is the number of residues and F the number of frequencies. Columns are standardised to zero mean and unit variance prior to analysis. Principal component analysis is applied independently to each dataset.

Mahalanobis distance in PCA space

Let $\mathbf{Z}$ denote the matrix of PCA scores. The centroid $\boldsymbol{\mu}$ and covariance matrix $\boldsymbol{\Sigma}$ of $\mathbf{Z}$ are computed and the Mahalanobis distance for each residue n is defined as:

$$D_M(n) = \sqrt{(\mathbf{z}_n - \boldsymbol{\mu})^\top \boldsymbol{\Sigma}^{-1} (\mathbf{z}_n - \boldsymbol{\mu})} \tag{18}$$

The maximum observed distance is retained as a dataset-level Mahalanobis score.

Q-Score

The standardised data are reconstructed from the PCA model and compared to the original standardised matrix. For each residue n, the reconstruction error is computed as:

$$Q(n) = \|\mathbf{x}_n - \hat{\mathbf{x}}_n\|^2 \tag{19}$$

Q-scores are standardised, and the fraction of residues exceeding the 90th percentile of the Q-score distribution is recorded as a dataset-level metric.

Aggregate Score

For each dataset, multiple complementary consistency metrics are retained, capturing distribution-level agreement, multivariate coherence and residue-level variability across magnetic fields. The retained metrics are:

- Kolmogorov–Smirnov statistic (KS)
- Inverted quantile–quantile agreement score (Q–Q $R^2$)
- Median absolute deviation of fractional changes ($MAD_Q$)
- Mahalanobis distance–based multivariate deviation score
- Fraction of residues exceeding the 90th percentile of the PCA-based Q-score distribution

Each metric is rescaled to the [0,1] interval using min–max normalisation. A single dataset-level inconsistency score is then defined as a weighted sum of the rescaled metrics.

The total inconsistency score for dataset d, denoted S(d), is computed as:

$$S(d) = w_{Ks}KS(d) + w_{QQ}(QQ - R^2)(d) + w_{MADQ}MAD_Q(d) + w_{Mah}Mah'(d) \quad (20)$$

with fixed weights:

$$\begin{aligned} w_{\mathrm{KS}} &= 0.30 \\ w_{\mathrm{QQ}} &= 0.40 \\ w_{\mathrm{MADQ}} &= 0.05 \\ w_{\mathrm{Mah}} &= 0.15 \end{aligned}$$

The chosen weights reflect the relative statistical stability, scope of information captured and sensitivity of each metric to systematic discrepancies across magnetic fields, as observed empirically across the analysed datasets.

The inverted Q–Q $R^2$ metric is assigned the largest weight, as it evaluates agreement between entire empirical distributions of J(0) values using matched quantiles. This measure is robust to outliers, insensitive to absolute scaling and sensitive to systematic shifts, changes in spread and non-linear distortions between datasets, providing a stable global indicator of field-to-field agreement.

The KS statistic contributes substantially as a complementary distribution-level measure that captures the maximum deviation between cumulative distributions. While sensitive to localised discrepancies, it can be influenced by sample size and tail behaviour; its weighting balances sensitivity with robustness when combined with the Q–Q metric.

The Mahalanobis distance score quantifies multivariate deviations across all magnetic fields simultaneously in PCA space, enabling detection of residues exhibiting inconsistent behaviour in a joint-field context. Its moderate weight reflects its utility for multi-field datasets while limiting sensitivity to covariance estimation and extreme outliers.

The fractional-change MADQ metric summarises residue-level dispersion in J(0) differences using a robust statistic. Because this metric is sensitive to small denominators and low J(0) values, it is assigned a low weight and serves primarily as a weak regularising term penalising widespread residue-level instability.

Together, this weighting scheme prioritises global distributional agreement, incorporates multivariate coherence across magnetic fields and retains residue-level heterogeneity information without allowing any single metric to dominate the overall score. The weights were selected to ensure numerical stability, interpretability and robustness across a heterogeneous collection of BMRB-derived datasets, enabling objective identification of datasets suitable for reliable validation and benchmarking of ModA.

### Additional Supplementary Tables

| Category | Experiment | Timescale | Physical process probed | Primary observables | Typical extracted parameters | Modelling |
|---|---|---|---|---|---|---|
| **Fast internal motion** | $T_1$ ($R_1$) | ps–ns | Longitudinal spin–lattice relaxation | $R_1$ relaxation rates | Spectral density J(ω) order parameters $S^2$ local correlation times | Redfield theory spectral density models model-free (Lipari–Szabo) |
| **Fast internal motion** | $T_2$ ($R_2$) | ps–ns | Transverse spin–spin relaxation | $R_2$ relaxation rates | Spectral density J(ω) order parameters $S^2$ | Redfield theory spectral density models model-free (Lipari–Szabo) |
| **Fast internal motion** | Heteronuclear NOE | ps–ns | High-frequency internal motion | {1H}-X NOE enhancements | Order parameters $S^2$ fast motional amplitudes | Redfield theory spectral density models model-free (Lipari–Szabo) |
| **Fast internal motion** | Cross-correlated relaxation (CCR) | ps–ns | Interference between relaxation mechanisms | CCR rates η | Relative tensor orientations correlated motion | Redfield theory cross-correlation models |
| **Relaxation Dispersion** | CPMG relaxation dispersion | μs–ms | Exchange-induced transverse relaxation | $R_{2eff}$ vs νCPMG | Exchange rates kex populations chemical shift differences Δω | Bloch–McConnell relaxation-dispersion models |
| **Relaxation Dispersion** | $R_{1\rho}$ relaxation dispersion | μs–ms | Exchange under spin-lock field | $R_{1\rho}$ vs spin-lock power | Exchange rates kex populations Δω | Bloch–McConnell spin-lock dispersion models |
| **Relaxation Dispersion** | Exchange-induced line broadening ($\Delta R_2$) | μs–ms | Lifetime broadening due to exchange | $\Delta R_2$ contributions | kex model-dependent populations | Bloch–McConnell lifetime-broadening models |
| **Slow chemical exchange** | CEST | ms–s | Saturation transfer between exchanging states | Intensity vs saturation offset | Exchange rates minor-state populations Δω | Bloch–McConnell saturation-transfer models |
| **Slow chemical exchange** | EXSY | ms–s | Magnetization transfer during mixing time | Cross-peak intensities vs τmix | kAB kBA state populations | Bloch–McConnell exchange kinetics |
| **Slow chemical exchange** | ZZ exchange | ms–s | Longitudinal population exchange | Diagonal and exchange peak intensities | kAB kBA populations | Bloch–McConnell longitudinal exchange models |
| **Slow chemical exchange** | DEST | ms–s | Exchange with NMR-invisible states | Signal attenuation vs saturation | Exchange rates populations bound-state $R_2$ | Bloch–McConnell dark-state exchange models |

**Supplementary Table 5. NMR dynamics experiments.** Summary of representative NMR relaxation and exchange experiments grouped by the timescale and physical process probed. For each experiment, the table lists the primary experimental observables, typical extracted dynamic or kinetic parameters and the corresponding modelling framework, spanning fast ps-ns internal motion, µs-ms relaxation dispersion and slower ms-s chemical exchange.

| Package | Category | $R_1$ | $R_2$ | hetNOE | Lipari-Szabo | CPMG | R1ρ relaxation dispersion | CEST | DEST | EXSY | ZZ | Etas | Multi-purpose Multi-Fitting | API - Scripting | Interactive analysis to Raw Data | Plugins |
|---|---|---|---|---|---|---|---|---|---|---|---|---|---|---|---|---|
| CcpNmr[1] | Academic | ✓ | ✓ | ✓ | ✓ | ✓ | ✓ | ✓ | ✓ | ✓ | ✓ | ✓ | ✓ | ✓ | ✓ | ✓ |
| Relax[15] | Academic | ✓ | ✓ | ✓ | ✓ | ✓ | ✓ | | | | | | | ✓ | | |
| ModelFree[14] | Academic | ✓ | ✓ | ✓ | ✓ | | | | | | | | | ✓ | | |
| TENSOR2[26] | Academic | ✓ | ✓ | ✓ | ✓ | | | | | | | | | | | |
| NESSY[27] | Academic | | | | | ✓ | ✓ | | | | | | | | | |
| ChemEx[28] | Academic | | | | ✓ | ✓ | ✓ | ✓ | ✓ | | | | ✓ | ✓ | | |
| GUARDD[29] | Academic | | | | | ✓ | | | | | | | | | | |
| CPMGFit | Academic | | | | | ✓ | | | | | | | | | | |
| CATIA | Academic | | | | | ✓ | | | | | | | | | | |
| ShereKhan | Academic | | | | | ✓ | | | | | | | | | | |
| GLOVE | Academic | | | | | ✓ | | | | | | | | | | |
| RING NMR[30] | Academic | ✓ | ✓ | ✓ | | ✓ | ✓ | ✓ | | | | | ✓ | | ✓ | |
| DESTfit | Academic | | | | | | | | ✓ | | | | | | | |
| EXSYCalc | Academic | | | | | | | | | ✓ | | | | | | |
| NMRFx[31] | Academic | | | | | | | | | | | | | ✓ | ✓ | ✓ |
| RotDiff[32] | Academic | | | | ✓ | | | | | | | | | | | |
| Top Spin[33] | Vendor / Commercial | ✓ | ✓ | ✓ | | | | | | ✓ | ✓ | | | ✓ | | ✓ |
| Mnova[34] | Vendor / Commercial | ✓ | ✓ | ✓ | | | | | | | | | | ✓ | ✓ | ✓ |
| Agilent | Vendor / Commercial | | | | | | | | | | | | | ✓ | | ✓ |
| Jeol-jason | Vendor / Commercial | ✓ | ✓ | ✓ | | | | | | ✓ | ✓ | | | ✓ | ✓ | ✓ |

**Supplementary Table 6. Comparison of NMR dynamics analysis software.** Summary of representative software packages used for biomolecular NMR relaxation and exchange analysis. Features are grouped by supported experiment types, modelling capabilities, scripting or API access, interactive linkage to experimental data and extensibility through plugins. Check marks indicate whether each package provides native or documented support for the corresponding capability.

| Name | Primary Experiment Tags | Category | Language | Github |
|---|---|---|---|---|
| **CorrFunction_NMRRelaxation** | $R_1/T_1$; $R_2/T_2$; heterNOE (from MD correlation functions) | Notebook/Repo | Python | 1 |
| modelfree-protein15n | Model-free ($^{15}$N $R_1/R_2$/ heterNOE) | Package (PyPI) | Python | 2 |
| relaxometrynmr | $T_1$; T1rho; $T_2$ (relaxometry fitting) | Package (PyPI) | Python | 3 |
| relaxNMR | $^1$H TD relaxometry; ILT | Package/Repo | Python | 4 |
| pyParaTools | PRE (paramagnetic relaxation enhancement); PCS; RDC fitting/analysis | Package/Repo | Python | 5 |
| BMNS (Bloch-McConnell N-State) | $R_{1\rho}$ dispersion; (also $R_{2eff}$ fits per docs) | Package/Tool | Python | 6 |
| xpy3-tools | TopSpin Python scripting (processing/automation; may include series handling) | Script collection/Repo | Python | 7 |
| TRACT_analysis (nomadiq) | TRACT | Script/Repo | Python | 8 |
| NMRforMD (nmrformd) | $T_1/T_2$ from MD trajectories | Script/Repo | Python | 9 |
| papuaNMR_Exchange (nomadiq) | Nz exchange / EXSY-style exchange fitting | Tool/Repo | Python | 10 |
| MERT-NMR | Multidimensional time-domain relaxation parameter estimation | Package/Repo | MATLAB | 11 |
| BM_sim_fit (cest-sources) | Bloch-McConnell simulation+fit (CEST) | Tool/Repo | MATLAB | 12 |
| NMRAnalysis.jl | Relaxation (incl $T_1/T_2$); diffusion; TRACT | Package | Julia | 13 |
| NMRFx Rate Analysis (docs) | General rate analysis panel (intensity→rates; fits) | Suite/docs | Java-based suite | 14 |
| eMF (easy Model-Free) | Model-free ($^{15}$N relaxation) | Tool/Repo | C++/Python | 15 |
| OpenVnmrJ source (legacy vendor ecosystem) | Vendor macro-based relaxation/exchange workflows (legacy) | Suite (vendor ecosystem) | C/Java/etc | 16 |
| MINOTAUR | High-resolution relaxometry + high-field rates (shuttling) | Program/Repo | C / Python | 17 |
| SpinRelax (zharmad) | Spin relaxation parameter computation + fit-to-experiment (MD-based) | Workflow/Repo | Bash/Python | 18 |
| neural-fitting (momentum-laboratory) | Bloch-McConnell ODE fitting via neural methods (CEST/exchange) | Research code/Repo | Python | 19 |
| nmrformd | MD NMR | Research code/Repo | Python | 20 |
| comdnmr | Ring repo | Program/Repo | Java-based suite | 21 |
| Rela2x | Relaxation Theory | Research code/Repo | Python | 22 |
| BD-NMR | Brownian Dynamics NMR | Research code/Repo | C++/Cuda | 23 |
| flint | CPMG relaxation dispersion/ $R_1\rho$ dispersion | Program/Repo | MATLAB/Python | 24 |
| idpemery/nmr-relaxation | various scripts | Tool/Repo | MATLAB/Python | 25 |
| IDP Dynamics Project | intrinsically disordered proteins | Tool/Repo | Jupyter /Python | 26 |
| NMR_T1_fits | time-resolved $T_1$ relaxation constants from spoiler saturation recovery NMR | Tool/Repo | Python | 27 |
| GUARDD | rigorous analysis of CPMG RD NMR data. | Tool/Repo | MATLAB | 28 |
| PyRelax | various scripts for Redfield theory | Tool/Repo | Python | 29 |
| cifit | Legacy repo for cifit program | Program/Repo | c | 30 |

**Supplementary Table 7. GitHub-hosted scripts and repositories for NMR dynamics analysis.** Summary of publicly available GitHub-hosted scripts, notebooks and repositories relevant to biomolecular NMR relaxation or dynamics analysis. Entries are annotated by primary experiment tags, repository category, implementation language and GitHub availability, highlighting the diversity of community-developed tools and lightweight analysis resources outside larger integrated software packages.

https://github.com/achicks15/CorrFunction_NMRRelaxation https://pypi.org/project/modelfree-protein15n/ https://pypi.org/project/relaxometrynmr/ https://github.com/specmicp/relaxNMR
https://github.com/mscook/pyParaTools
https://github.com/IsaacJK/BMNS
https://github.com/olegvpetrov/xpy3-tools https://github.com/nomadiq/TRACT_analysis https://github.com/simongravelle/nmrformd https://github.com/nomadiq/papuaNMR_Exchange https://github.com/genematx/MERT-NMR
https://github.com/cest-sources/BM_sim_fit https://github.com/waudbygroup/NMRAnalysis.jl https://github.com/onemoonsci/nmrfxprocessordocs/blob/master/pages/02.viewer/06.ratesandbinding/01.rateanalysis/docs.md https://github.com/sunghunbae/eMF
https://github.com/OpenVnmrJ/OpenVnmrJ https://github.com/nbolikcoulon/MINOTAUR https://github.com/zharmad/SpinRelax
https://github.com/momentum-laboratory/neural-fitting https://github.com/simongravelle/nmrformd https://github.com/brucejohnson/comdnmr https://github.com/hillaper/Rela2x
https://github.com/mircozerbetto-unipd/BD-NMR https://github.com/paultnz/flint
https://github.com/idpemery/nmr-relaxation https://github.com/sdey17/IDP_Dynamics https://github.com/lfillbrook/NMR_T1_fits https://github.com/mpfosterlab/GUARDD
https://github.com/nbolikcoulon/PyRelax

https://github.com/ZryletTC/cifit

| Component | Feature | Implementation Summary | Rationale and impact |
|---|---|---|---|
| Architecture | Modular framework | Separates spectral-density theory, rotational diffusion, relaxation-rate calculation, optimisation, uncertainty estimation, and reporting into independent components. | Makes assumptions explicit, improves reproducibility, and allows individual components to be extended or replaced. |
| Architecture | Registry-based spectral-density models | Implements Lipari-Szabo model parameterisations as independent classes with explicit optimised parameters. | Enables new model forms to be added without altering diffusion, relaxation-rate, or optimisation code. |
| Diffusion modelling | Unified rotational diffusion framework | Implements isotropic, axially symmetric, and fully anisotropic tumbling models within a common workflow. | Enables systematic testing of how global tumbling assumptions affect residue-specific dynamic parameters. |
| Structural geometry | Tensor-based bond-vector handling | Uses PDB or mmCIF coordinates to define N-H bond-vector orientations and initialise anisotropic diffusion geometry. | Incorporates structural geometry into diffusion modelling without imposing a physical model of internal motion. |
| Initialisation | Robust initial $\tau_c$ estimation | Estimates initial global tumbling from filtered $R_1$ and $R_2$ data after excluding low-HETNOE or anomalous residues. | Reduces bias from flexible, exchanging, or poorly behaved residues before model-free fitting. |
| Initialisation | Data-derived initial order parameter | Derives an initial average order parameter from robust statistics of $R_1$ and $R_2$ products. | Provides physically informed starting values instead of arbitrary initial S2 estimates. |
| Initialisation | Adaptive residue-specific $\tau_e$ bounds | Adjusts tau-e bounds using $S^2$, $\tau_c$, $R_2/R_1$ rankings, optional hetNOE information, and exchange indicators. | Gives rigid and flexible residues appropriate search spaces and reduces parameter degeneracy. |
| Optimisation | Differential Evolution global search | Uses population-based Differential Evolution to fit nonlinear Lipari-Szabo parameter spaces. | Reduces dependence on initial guesses and improves recovery from local or correlated minima. |
| Optimisation | Reproducible optimisation presets | Provides low-, medium-, high-accuracy, and custom Differential Evolution configurations. | Balances speed and precision while retaining transparent and reproducible optimiser behaviour. |
| Optimisation | Expert optimiser configurability | Exposes strategy, population size, mutation, recombination, tolerance, initialisation, dithering, and polishing controls. | Permits controlled analysis of difficult datasets without relying on hidden optimiser defaults. |
| Objective function | Noise-adaptive chi-squared fitting | Treats $R_1$, $R_2$, and hetNOE as separate noise groups with adaptive residual models and stabilised uncertainty weighting. | Better reflects heterogeneous relaxation-data errors and reduces sensitivity to unreliable uncertainties. |
| Objective function | Robust outlier handling | Applies optional Huber-loss weighting to standardised residuals during fitting. | Limits the influence of outlying measurements while preserving least-squares behaviour for well-fit data. |
| Uncertainty estimation | Latin Hypercube Monte Carlo refitting | Generates stratified synthetic datasets and refits each through the full optimisation workflow. | Improves uncertainty-space coverage and captures nonlinear coupling between fitted parameters. |
| Uncertainty estimation | Physically constrained perturbation model | Samples Monte Carlo perturbations from truncated distributions with optional constraints such as positive rates. | Avoids unrealistic pseudo-datasets that can destabilise refitting or inflate uncertainty estimates. |
| Uncertainty estimation | Robust uncertainty statistics | Estimates final parameter uncertainties using robust statistics such as the median absolute deviation. | Reduces sensitivity to unstable refits and non-Gaussian parameter distributions. |
| Model selection | Penalised information-criterion scoring | Compares candidate models using AIC, BIC, AICc, or BICc scores. | Balances fit quality against model complexity and reduces overfitting in limited datasets. |
| Model selection | Monte Carlo model-confidence estimates | Recomputes model selection across pseudo-datasets and reports model frequencies as confidence values. | Distinguishes strongly supported residue models from statistically ambiguous assignments. |
| Global refinement | Post-fit global tau-c optimisation | Refines a single diffusion-scaling factor after residue-level fitting while keeping internal parameters fixed. | Enforces consistency between local fitted dynamics and global molecular tumbling. |
| Performance | Parallel residue-wise execution | Distributes independent residue-level model fitting and Monte Carlo refitting across worker processes. | Makes exhaustive model comparison and uncertainty analysis practical for larger datasets. |
| Workflow | CLI-driven reproducible analysis | Runs from explicit input and settings files that can also be wrapped by a graphical interface. | Combines scripted reproducibility with interactive usability in CcpNmr-based workflows. |
| Data integration | Flexible input and structural formats | Supports tabular relaxation data, saved settings, structural coordinates, and planned CSV, Excel, JSON, NEF, PDB, mmCIF, and CcpNmr inputs. | Reduces manual conversion and improves integration with existing NMR analysis pipelines. |
| Reporting | Transparent residue-level outputs | Reports fitted parameters, uncertainties, model scores, confidence values, and visual model-selection summaries. | Supports critical inspection of parameter reliability and model-selection confidence across the sequence. |

**Supplementary Table 8. ModelAnalysis implementation features.** Summary of the main ModA components for Lipari-Szabo analysis, including architecture, diffusion modelling, optimisation, uncertainty estimation, model selection, performance and reporting. The table lists each feature, its implementation and its rationale within the AnalysisDynamics workflow.
